# Survivin Mediates Mitotic Onset in HeLa Cells Through Activation of the Cdk1-Cdc25B Axis

**DOI:** 10.1101/2023.12.21.572935

**Authors:** Pedro M. Cánovas

**Author notes:** Correspondence: Pedro M. Cánovas, Ph.D., 38 Mount Guyot Street, North Brookfield, MA 01535.

## Abstract

The Survivin protein has roles in repairing incorrect microtubule-kinetochore attachments at prometaphase and the faithful execution of cytokinesis, both as part of the *<u>c</u>hromosomal <u>p</u>assenger <u>c</u>omplex* (CPC) (1). In this context, errors frequently lead to aneuploidy, polyploidy and cancer (1). Adding to these well-known roles of this protein, this paper now shows for the first time that Survivin is required for cancer cells to enter mitosis, and that, in its absence, HeLa cells accumulate at early prophase, or prior to reported before (2, 3). The early prophase blockage in cells lacking Survivin is demonstrated by the presence of an intact nuclear lamina and low Cdk1 activity (4). Interestingly, Survivin and Cdk1 form a complex *in vivo*. This interaction peaks at mitosis, and its molecular targeting indicates that Survivin is needed for Cdk1 to be active. In this regard, escaping the blockage induced by Survivin abrogation leads to multiple mitotic defects, or *mitotic catastrophe*, and eventually cell death. Mechanistically, recombinant Survivin can induce the activation of Cdk1 via Cdc25 *in vitro*. Coincidentally, Cdk1 mislocalizes at the centrosome when Survivin is not expressed. Moreover, Survivin directly interacts with phosphatase Cdc25B, both *in vitro* and *in vivo*, and in the absence of the former, an inactive cytosolic Cdc25B-Cdk1-Cyclin B1 complex accumulates, which coincides with the mitotic arrest induced by Survivin depletion. Finally, in agreement with a role for Survivin in the early activation of Cdk1, the G2/early prophase accumulation induced in HeLa cells by Survivin abrogation could be bypassed by a gain-of-function Cdc25B mutant, which drove cells into mitosis.

## INTRODUCTION

The sequential activation of the Cdk1 protein kinase is an essential requirement for correct entry of cells into mitosis (5). This process is known in detail, and involves formation of a complex between Cdk1 and its activator Cyclin B1 (5), and removal of inhibitory phosphates on residues Threonine 14 and Tyrosine 15 of Cdk1 by the phosphatase Cdc25 (5). On this latter step, Cdk1 can be activated by three Cdc25 isoforms (5), being generally accepted that Cdc25B is the first one that acts on the kinase at the centrosome (6, 7), and that the other two amplify the centrosomal signal in the cytosol and the nucleus (5).

Full activation of Cdk1 in the nucleus at late prophase is known to be a *point-of-no-return* at mitosis (8). Here, the nuclear lamina is pulled apart as a result of Cdk1 phosphorylation of the Lamin B subunits, triggering disassembly of the nuclear membrane (4, 9, 10). Following nuclear membrane disassembly, cytosolic microtubules are able to reach the kinetochores assembled on the centromeres, as cells transverse from late prophase to prometaphase (11, 12). Interestingly, this so-called *mitotic commitment*, which requires high levels of Cdk1 activity in the nucleus (13, 14), is a phenotype difficult to discern, and only a detailed microscopic and biochemical analysis of mitotic cells can differentiate between those that have irreversibly entered mitosis, and those that are not fully committed yet. In this context, a frail activation of Cdk1 often leads to *mitotic catastrophe* and cell death (15).

On a different aspect of mitosis, it is well-known the role of the Survivin protein in microtubule dynamics from prometaphase throughout cytokinesis (1). Here, Survivin malfunction leads to multiple mitotic defects (2, 3, 16, 17). Specifically, on the role of Survivin at prometaphase, this protein is part of the CPC together with Borealin, INCENP and Aurora B, where it contributes to proper localization and activation of the complex at the centromere (1). Once at this location, Aurora B, the CPC’s catalytic subunit, can sense kinetochore tension, and help correcting faulty microtubule-kinetochore attachments in partnership with the microtubule depolymerase MCAK (1). In this scenario, mistakes caused by CPC’s malfunction generally lead to a sustained spindle checkpoint that, when overridden, produces fatal consequences, as entire chromosomes may mis-segregate giving rise to aneuploidy, a typical feature of many tumors (1). On the other hand, during cytokinesis, lack of Survivin function often leads to formation of giant multinucleated cells through endoreplication (1).

Survivin has also been localized in the centrosome (18) but its role at this non-membranous organelle has not been fully elucidated yet. In this regard, Survivin depletion causes a *mini spindle* phenotype in *Xenopus* egg extracts (17) and HeLa cells (16, 19), which points at a failure in centrosome separation in actively-dividing cells, an early mitotic event for which centrosomal Cdk1 activity is essential through its effect on Eg5 (20, 21). These data, and the fact that Survivin binds to both Cdk1 (22) and microtubules (17, 23), seems to suggest a role of Survivin in the proper localization and/or activation of the main mitotic kinase at the centrosome. However, despite the interest of this topic, and its possible important implications in cancer development, the role of Survivin at an earlier time than prometaphase has not been studied. For this reason, with the objective to unravel the role of Survivin at mitotic onset in cancer cells, I decided to revisit the early mitotic events that follow abrogation of Survivin in HeLa cells.

## RESULTS

### siRNA-Mediated Loss of Survivin Induces an Early Prophase Blockage

To clarify the stage at which cancer cells first arrest at mitosis following Survivin abrogation, I first had to reproduce the G2/M-phase blockage reported by other authors in the absence of Survivin (2, 16). For that purpose, I needed a Survivin siRNA that could cause an acute and reproducible Survivin abrogation, and concomitant G2/M-phase arrest. It was reported before that the S4 Survivin siRNA can induce both these phenotypes (16). Therefore, I decided to use this oligonucleotide in my preliminary RNAi experiments. As Fig. 1A shows, transfection of asynchronous HeLa cell cultures with the Survivin siRNA (S4) caused a complete and reproducible ablation of this protein, as compared to the control siRNA (VIII).

**FIG. 1.**
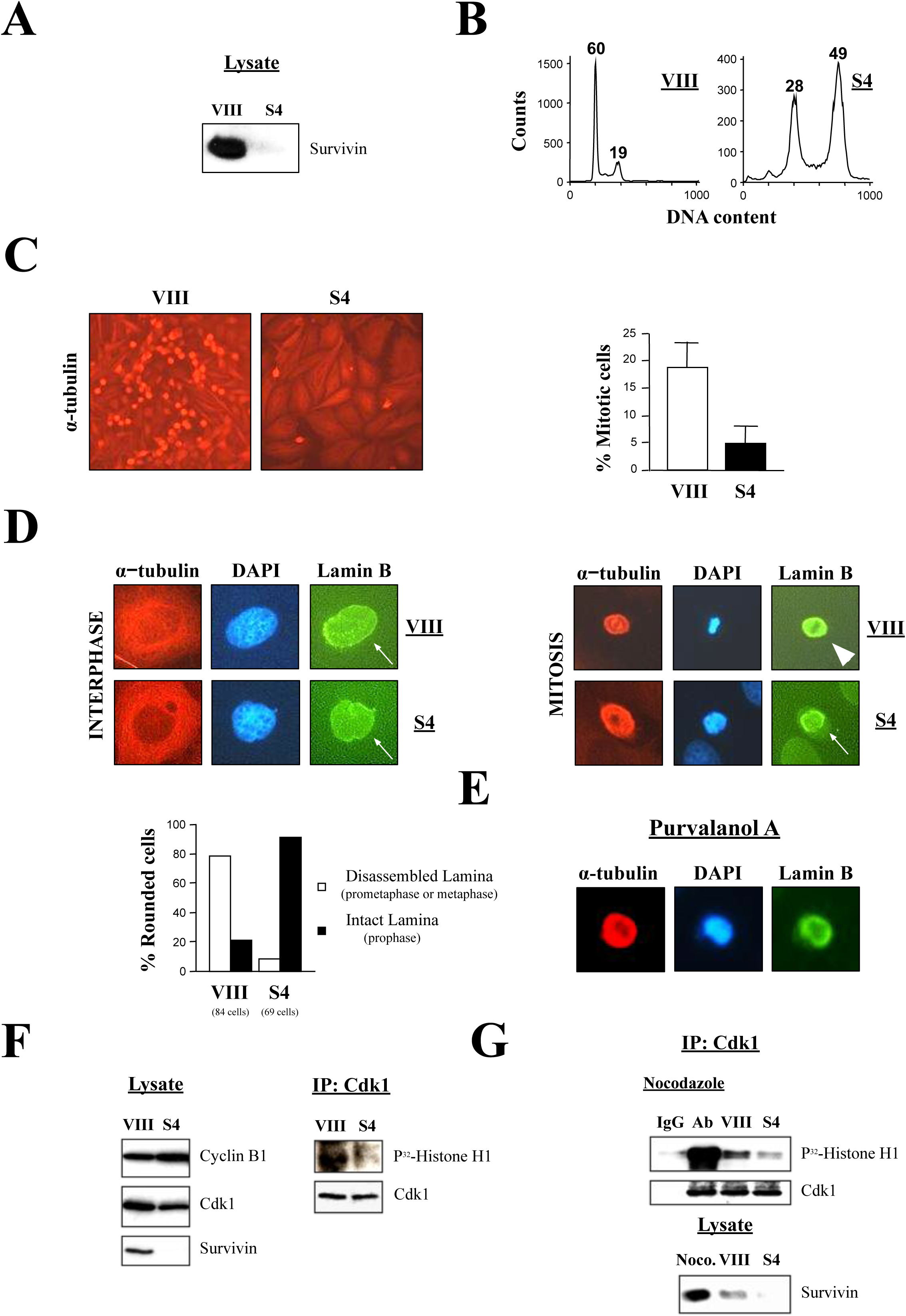
siRNA-mediated loss of Survivin induces an early prophase blockage in asynchronous HeLa cell cultures. *A*, *B*, Survivin knockdown. HeLa cells transfected with control (VIII) or Survivin (S4) siRNA were analyzed by Western blotting (*A*) or FACS analysis (*B*). The percentage of G1 (*2N*), G2/M-phase (*4N*) and polyploid (*>4N*) cells is indicated. *C*, siRNA-treated HeLa mitotic cells. HeLa cells transfected with the indicated siRNA were analyzed by fluorescence microscopy with an antibody to α-tubulin (*left*). Bar graph shows the percentage of mitotic cells in transfected cultures (n=4) (*right*). *D*, Nuclear lamina integrity in siRNA-treated HeLa cells. Interphase or mitotic HeLa cells transfected with the indicated siRNA, stained with an antibody to Lamin B or α-tubulin, and analyzed by fluorescence microscopy. DNA was stained with DAPI. Cells are shown at the same scale. Arrows show intact nuclear lamina, and arrow head points out at Lamin B colocalization with the mitotic spindle. The percentage of rounded mitotic cells with intact or disassembled nuclear lamina is indicated (*bottom*). Control (VIII) siRNA: 84 cells/8 fields; Survivin (S4) siRNA: 69 cells/12 fields (n=3). *E*, Purvalanol A treatment of HeLa cells. HeLa cells were treated with the Cdk1 inhibitor purvalanol A (20 μM), stained with an antibody to Lamin B or α-tubulin, and analyzed by fluorescence microscopy. DNA was stained with DAPI. *F*, Cdk1 kinase assay of siRNA-treated HeLa cells. siRNA-transfected HeLa cell lysates were used to immunoprecipitate (IP) Cdk1, and the kinase activity was analyzed by a Histone H1 phosphorylation assay (*right*). Amounts of Cyclin B1, Cdk1 and Survivin were analyzed by Western blotting as a control (*left*). *G*, Survivin ablation vs. prometaphase blockage. HeLa cells were treated with 10 μM nocodazole, and their Cdk1 activity was analyzed by an IP and a Histone H1 phosphorylation assay. Survivin siRNA-transfected HeLa cells (S4) were used as a control.

Next, I needed to check if the Survivin siRNA could induce a robust G2/M-phase blockage. To this end, I treated Hela cells with the control (VIII) or Survivin (S4) siRNA as in Fig. 1A, and subjected these samples to FACS analysis. As shown in Fig. 1B, HeLa cells not expressing Survivin but not control cells, accumulated at the G2/M-phase peak (28% of Survivin-depleted cells vs. 19% of control cells), or had a more than *4N* DNA content, as a result of endoreplication (49% of Survivin-depleted cells vs. 0% of control cells). Notice that in this experiment, no G1 cells (*2N* DNA content) could be seen in the absence of Survivin, indicating that no cells could complete mitosis.

Following the above preliminary experiments, I wanted to see the Survivin-depleted cells that were accumulating in the G2/M-phase peak (Fig. 1B) in order to determine where the Survivin-induced G2/M-phase blockage first occurred. To answer this question, I treated asynchronous HeLa cell cultures as before, and labeled these cells with an antibody against the α-tubulin protein, so I could analyze them by immunofluorescence microscopy. As Fig. 1C, *left*, VIII panel shows, control cultures contained many mitotic cells, as judged by the large number of rounded (i.e. prophase through anaphase) and splitting (i.e. telophase/cytokinesis) cells with bright microtubules that could be seen. In contrast to the control samples, HeLa cells where Survivin was ablated did not accumulate to a large extent in mitosis, as concluded by the small number of rounded cells, and the absence of telophase figures in the pictures (Fig. 1C, *left*, S4 panel). Instead, many large multinucleated cells could be detected in the cells that lacked Survivin, in agreement with the FACS analysis (Fig. 1B). To accurately determine the extent of the defect in mitotic entry in the cells not expressing Survivin, I counted the number of rounded cells from several microscope fields (from now on, I did not analyze the cells in telophase-cytokinesis since these cells were almost absent in the Survivin-depleted samples). These results are summarized in Fig. 1C, *right*, and show that HeLa cell cultures where Survivin was depleted, contained about 30% less rounded cells than the control ones (5% vs. 18%).

The drop in mitotic cells in cultures where Survivin was ablated was also confirmed in siRNA-transfected HeLa cultures analyzed by immunofluorescence microscopy using an antibody against the *bona fide* mitotic protein Cyclin B1 (Fig. S1). In these images, I could see that, following Survivin abrogation, most cells had a dispersed Cyclin B1 protein throughout the cytosol, a typical phenotype of late G2 or early prophase cells (24). Also, the Survivin-depleted cells that managed to enter mitosis appeared to be mostly in prophase, as judged by the colocalization of their Cyclin B1 protein and DNA (24). In contrast to the cells devoid of Survivin, Cyclin B1 co-localized with the spindle in the control samples, a typical feature of cells in prometaphase or metaphase (25). Therefore, from these data, I concluded that abrogation of Survivin in HeLa cells leads to an impairment in the capacity of these cultures to commit to mitosis (i.e. prophase to prometaphase transition) (8).

In order to obtain better visual data on the Survivin-depleted mitotic cells detected in Fig, 1C, siRNA-treated rounded HeLa cells were further analyzed by immunofluorescence microscopy, this time at a higher resolution (Fig. 1D). Here, three cellular markers were used to distinguish between the different cell cycle phases, namely: i) microtubule length, as shown by an α-tubulin antibody, where short bright microtubules indicate cells in mitosis, ii) chromatin condensation, observed by DAPI staining, where condensed chromatin denotes mitosis cells from late prophase onwards, and iii) nuclear lamina integrity, as shown by a Lamin B antibody, which discriminates between interphase through prophase cells (i.e. intact nuclear lamina), and prometaphase and onwards (i.e. disassembled lamina) (4, 9, 10). The results of the above experiments are shown in Fig. 1D. First, the left panels indicate that staining of interphase cultures was the same, independently of the siRNA treatment, and consisted of cells with long microtubules and an intact nuclear lamina (arrows) encircling the chromatin. Second, most control rounded cells assembled a spindle containing condensed chromatin at the spindle equator, and had a disassembled nuclear lamina, a phenotype previously ascribed to nuclear Cdk1 activity (10), where the Lamin B monomers co-localized with the spindle (Fig. 1D, *right*, arrow head), also a typical mitotic feature (26). In contrast to the control samples, most of the Survivin-knocked down rounded cells had a partial condensed chromatin surrounded by layers of short microtubules that encircled the DNA, and an interphase-looking nuclear lamina (i.e. circular lamina), which also surrounded the chromatin (Fig. 1D, *right*, bottom panels). This result was indicative of a partially activated Cdk1 in the cells lacking Survivin (13, 14), and confirmed the prophase blockage seen before at lower resolution, however contradicted the previous data, which claimed a prometaphase arrest under similar conditions (2, 3). To determine the extent of the intact lamina phenotype observed in the cells lacking Survivin, a number of siRNA-treated rounded cells from several independent experiments were scored for the presence of a Lamin B ring surrounding their chromatin. Fig. 1D, *bottom* shows that while 80% of the rounded cells treated with the control siRNA (VIII) had a disassembled lamina (i.e. cells in prometaphase and onwards), only 10% of the Survivin-depleted cells had a similar phenotype. If a role for Survivin in the full activation of Cdk1 at prophase were true, HeLa cells treated with a Cdk1 inhibitor should show a similar phenotype to the Lamin B ring observed when Survivin was absent. To test this possibility, HeLa cells were treated with the Cdk1 inhibitor purvalanol A at a concentration 20 μM, and samples were analyzed by immunofluorescence microscopy as above. As shown in Fig. 1E, the purvalanol A treatment replicated the intact-lamina and partially-condensed-chromatin phenotype observed in Survivin-depleted cells, further supporting a role for the Survivin protein in Cdk1 activation at prophase.

To obtain direct evidence of the Cdk1 function being affected by Survivin expression, I decided to check the kinase’s activity in the presence and absence of Survivin. To this aim, siRNA-transfected HeLa cultures were lysed, subjected to Cdk1 immunoprecipitation (IP), and their kinase activity was measured by a Histone H1 phosphorylation assay. The results of these experiments are presented in Fig. 1F, *right*, and show that the activity of the Cdk1 protein was reduced in the Survivin-depleted cells in comparison to the control. Also, the same lysates did not show major differences in the amounts of Cdk1 and Cyclin B1 (Fig. 1F, *left*), demonstrating that the lower Cdk1 activity in the Survivin-depleted HeLa cells was not due to a reduction in the amount of the proteins contributing to the Cdk1-Cyclin B1 complex. The Histone H1 phosphorylation data again clearly opposed the previous findings, which claimed an accumulation of cells at prometaphase following Survivin abrogation (2, 3).

To clearly show the difference between the blockage caused by Survivin abrogation, and a real high Cdk1 activity prometaphase arrest (27, 28), HeLa cells were either treated with the control (VIII) or Survivin (S4) siRNA as before, or incubated with the microtubule-depolymerizing agent nocodazole, which causes a potent prometaphase blockage, and their Cdk1 activities were compared by using a Histone H1 phosphorylation assay. The result of this experiment is shown in Fig. 1G. Here, it can be seen that HeLa cells lacking Survivin had a low Cdk1 activity, as already shown, which contrasted with the potent Cdk1 activation observed when similar cells were incubated with nocodazole. From these data and the rest of the results above, I concluded that HeLa cells block at early prophase with low Cdk1 activity in the absence of Survivin.

### The Early Prophase Blockage Caused by Survivin Abrogation in Asynchronous Cultures Can Be Replicated in Synchronized HeLa Cells, Precedes Endoreplication and Can Be Rescued by Exogenous Survivin

In order to pinpoint when absence of Survivin first impairs mitotic progression, I decided to repeat the above RNAi experiments, this time using synchronous HeLa cell cultures. Fig. 2A, *bottom* shows the FACS analysis of control (VIII) or Survivin (S4) siRNA-transfected, synchronized HeLa cells (Fig. 2A, *top* shows the Survivin content in the synchronized transfected cells). As it can be seen in this experiment, for the first 9 h, HeLa cells behaved exactly the same, and smoothly transitioned from G1 to G2/M-phase, independently of the siRNA treatment, and only behaved a bit sluggish, probably due to the siRNA transfection (Fig. 2A, *bottom*). The first divergence between the differently transfected cells appeared 14 h into the time course. Here, control HeLa cells started coming out of mitosis (i.e. G1 peak increase), and continued this trend for the remainder of the experiment (Fig. 2A, *bottom*, upper graphs). In contrast, Survivin-depleted cells remained arrested, or minimally escaped the G2/M-phase blockage and endoreplicated (Fig. 2A, *bottom*, lower graphs).

**FIG. 2.**
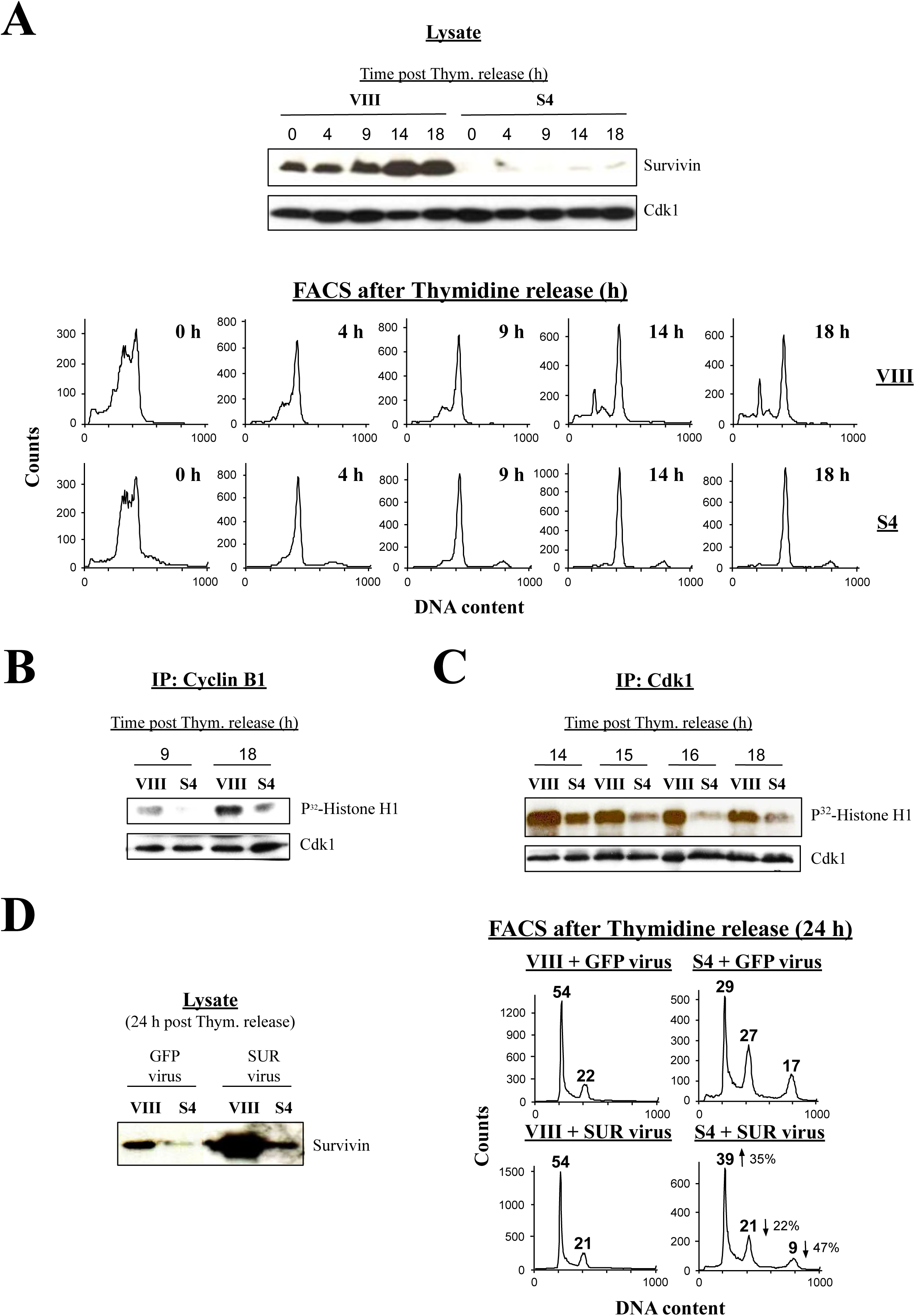
Survivin abrogation and reintroduction in synchronous HeLa cell cultures partially rescues the Survivin-induced blockage. *A*, Cell cycle analysis of siRNA-treated synchronized HeLa cells. HeLa cells transfected with control (VIII) or Survivin (S4) siRNA were synchronized with 2 mM thymidine for 48 h, released into fresh medium, and harvested at the indicated time intervals. Collected samples were subjected to Western blotting (*top*) or FACS analysis (*bottom*). *B*, *C*, Cdk1 activity during mitotic transition in siRNA-treated HeLa cells. Synchronized HeLa cells, previously transfected with the indicated siRNA, were harvested at the indicated time points following their release. Lysates were prepared and used to immunoprecipitate (IP) Cyclin B1 (*B*) or Cdk1 (*C*), and the immune complexes were analyzed in a Histone H1 phosphorylation assay. *D*, Rescue of G2/M-phase blockage induced by Survivin ablation. Synchronized HeLa cells transfected with the indicated siRNA were transduced, following the thymidine release, with an adenovirus encoding GFP (pAd-GFP) or GFP-Survivin (pAd-Survivin), and subjected to FACS analysis after 24 h (*right*). The percentage of G1 (*2N*), G2/M-phase (*4N*) and polyploid (*>4N*) cells is indicated. Arrows and percentages indicate changes in cell populations. Lysate panel shows expression of pAd-GFP or pAd-Survivin in siRNA-transfected HeLa cells (*left*). Experiment was repeated two more times with similar results.

To check whether synchronous HeLa cells depleted of Survivin were also impaired in Cdk1 activation as the asynchronous ones, and, if that were the case, determine when this anomaly first occurred, the Cdk1 activity of siRNA-transfected, synchronized HeLa cells was analyzed by a Histone H1 phosphorylation assay. Fig. 2B shows a Cyclin B1 IP, which also pulled Cdk1 down, of control and Survivin-depleted cells. Here, it can be seen that the samples lacking Survivin did not have any Cdk1 activity at 9 h, a time at which these cells approached the G2/M-phase checkpoint (see Fig. 2A, *bottom*), and very little kinase activity later in the time course (18 h). In contrast, as expected, Cdk1 activity gradually increased as control cells progressed through mitosis (Fig. 2B). To confirm this result, and find out in more detail when the Cdk1 kinase was first impaired in mitosis following Survivin abrogation, I performed a second IP, this time directly against the Cdk1 kinase, and using lysates of siRNA-transfected cultures, which were collected between 14 and 18 h post-thymidine release, or exactly when the control cells started exiting mitosis (see Fig. 2A, *bottom*, top graphs). As Fig. 2C shows, Survivin-knocked down cells had a low Histone H1 signal at all time points collected. In contrast, the control cells had a higher Cdk1 activity at the beginning of the time course (14 h), and, as expected, a gradual decrease, as cells continued exiting mitosis (18 h). From the Histone H1 results, I concluded that Survivin is required to activate Cdk1, and that in its absence, a strong G2/M-phase blockage ensues in HeLa cells.

The above results strongly connected the transition from G2/M-phase to G1 in Hela cells with Survivin expression and Cdk1 activity. To further prove this theory, re-expression of Survivin in samples where this protein had previously been ablated should push the G2/M-phase cells out of the blockage. To check this possibility, siRNA-transfected, synchronized HeLa cells were infected with a replication-deficient adenovirus encoding GFP (GFP virus) or GFP-Survivin (SUR virus) at the time of thymidine release, and lysates were analyzed by Western blotting (Fig. 2D, *left*) or FACS (Fig. 2D, *right*) 24 h later. As it can be seen, lack of Survivin followed by transfection with GFP virus did not rescue the G2/M-phase cell blockage (Fig. 2D, *right*, S4 + GFP virus). Notice here, the small G1 peak and the larger endoreplicated population in HeLa cells lacking Survivin as compared to Fig. 2A, *bottom*, lower graphs, which could routinely be seen in Survivin-depleted cells at very late time points (24 h) in the RNAi experiments. A plausible explanation for the G1 peak in the cells lacking Survivin might be some kind of recovery mediated by a small accumulation of Survivin at late time points (see Fig. 2D, *left*). On the other hand, the larger endoreplication peak could also be assigned to adaptation (i.e. *mitotic slippage*) at longer time courses. In contrast to the GFP virus control, expression of SUR virus in Survivin (S4) siRNA-transfected, synchronous HeLa cultures reduced 47% the population of endoreplicated cells and 22% the G2/M-phase blockage, and increased 35% the population of cells that completed cytokinesis and entered G1 (Fig. 2D, *right*, S4 + SUR virus, arrows). Here, it should be noticed that the Survivin siRNA-treated cells transfected with SUR virus showed a much lower adenoviral Survivin expression than the control cells, probably due to the still presence of the Survivin oligonucleotide in these samples, which most probably counteracted the effect of the exogenous Survivin protein on rescuing the Survivin-depleted cells.

### A Survivin Peptide Spanning Ala55 through Asp70 (SUR A55-D70) Binds the Cdk1 αC/β4 Loop

It is well known that the Survivin and Cdk1 proteins form a complex *in vivo* (22). Here, I confirmed this result, and showed that this interaction peaks as HeLa cells enter mitosis (Fig. S2A). To see whether the interaction between Survivin and Cdk1 is direct, I expressed and purified GST-Survivin and His-Cdk1 from *E. coli*, and used these recombinant proteins to carry out a GST-pull down experiment. As Fig. S2B shows, GST-Survivin but not GST alone could pull down the His-Cdk1 protein kinase *in vitro*, demonstrating that these two proteins bind directly to each other. As a consequence of this result, I could now address the question of whether Survivin is required for the direct activation of Cdk1 *in vivo* by making a peptide that could interfere with the cellular Survivin-Cdk1 complex. To this end, N- and C-terminal Survivin deletion mutants attached to a GST-tag were made, and used in GST-pull downs with His-Cdk1. Fig. S2C, *left* shows that His-Cdk1 interacted with the Survivin N-terminus (i.e. first 70 amino acids) but not the C-terminus (i.e. amino acids from 71 to 142). To narrow down the Cdk1-binding site in Survivin, more GST-Survivin deletion mutants, this time spanning residues 15 to 142 (GST-SUR 15-142), 38 to 142 (GST-SUR 38-142), 55 to 142 (GST-SUR 55-142) or 81 to 142 (GST-SUR 81-142) were generated. Fig. S2C, *right* shows the results of the GST-pull downs using these other Survivin fragments. From these data, it was concluded that the minimal region in Survivin that binds to Cdk1 comprises the residues from Ala55 (A55) to Asp70 (D70), a region in the BIR domain with an abundant number of acidic amino acids (29) (Figs. S3A and C).

Apart from the sequence in the Survivin protein contributing to the Survivin-Cdk1 interaction, I also wanted to know the region in the Cdk1 kinase that binds to Survivin. For this reason, I made His-Cdk1 deletion mutants containing residues 1 to 183 (His-Cdk1 1-183), 182 to 297 (His-Cdk1 182-297), 1 to 85 (His-Cdk1 1-85), 1 to 56 (His-Cdk1 1-56), 1 to 43 (His-Cdk1 1-43), 9 to 85 (His-Cdk1 9-85) or 36 to 85 (His-Cdk1 36-85) (Fig. S2D), and used them in binding assays with Survivin. Fig. S2D, *top* shows GST-Survivin pull downs using the Cdk1 mutants His-Cdk1 1-183, His-Cdk1 182-297 and His-Cdk1 1-85. These experiments indicated that only the Cdk1 fragments His-Cdk1 1-183 and His-Cdk1 1-85, which contain the kinase N-terminal region, could bind to GST-Survivin but not GST. Moreover, I also made and biotinylated the smallest Survivin region that binds to Cdk1 (SUR A55-D70), and used it in streptavidin-binding assays with all the His-Cdk1 deletion mutants (Fig. S2D, *bottom*). As seen, all the Cdk1 mutants but His-Cdk1 182-297, His-Cdk1 1-56 and His-Cdk1 1-43 bound to SUR A55-D70. From these data, I concluded that the region in Cdk1 that binds to Survivin is located in the kinase N-terminal sequence, and comprises the amino acids from Lys56 (K56) to Met85 (M85), or a sequence that includes the αC/β4 loop of the kinase (Figs. S3B and D).

### The SUR A55-D70 Peptide Targets the Survivin-Cdk1 Complex, Inducing its Disassembly, Loss of Cdk1 Activity, Mitotic Abnormalities and Apoptosis

Before analyzing the efficacy of the SUR A55-D70 peptide at interfering with the assembly of the Survivin-Cdk1 complex, and its effect on Cdk1 activity *in vivo*, I decided to carry out a few preliminary control experiments. First, I wanted to know whether the SUR A55-D70 peptide could pull down the Cdk1 protein from HeLa cell lysates. For this purpose, I incubated streptavidin beads pre-bound to the biotinylated scrambled or SUR A55-D70 peptide with HeLa cell extracts, and analyzed whether the endogenous Cdk1 protein bound or not to the resin. As Fig. S4A shows, endogenous Cdk1 bound to the beads that contained the Survivin peptide but not to the control reagent.

Second, I wanted to know whether the SUR A55-D70 peptide could displace the endogenous Survivin protein from its complex with Cdk1, and, if that was the case, whether this displacement affected the kinase activity. To that aim, I used a cell-free system, consisting of HeLa cell lysates from nocodazole-treated cultures, which should contain a large number of active Survivin-Cdk1 complexes. These cell extracts were supplemented with an ATP-regenerating system to make them biochemically active, and then incubated with the control or Survivin peptide for 1 h at 30°C. Fig. S4B shows that when the SUR A55-D70 peptide was incubated with nocodazole-treated HeLa cell lysates, and then Cdk1 was immunoprecipitated (IP), the kinase did not bind to the endogenous Survivin protein, and this correlated with a low Cdk1 activity in a Histone H1 phosphorylation assay. In contrast, no effect on Survivin binding to Cdk1 or the kinase activity was observed with the control peptide. These results clearly demonstrated that Cdk1 activity depends on its binding to the Survivin protein.

Finally, before using the SUR A55-D70 peptide to analyze its *in vivo* effect, I wanted to know whether this reagent could transverse the HeLa cell membrane when attached to an N-terminal HIV tat cell-permeable sequence. Fig. S4C shows that both the scrambled and Survivin A55-D70 peptides attached to the HIV sequence, readily accumulated inside the cells following their incubation with HeLa cultures.

After the above controls, asynchronous HeLa cultures were incubated with HIV tat-peptides (50 μM) for 6 h, and samples were analyzed by immunofluorescence using an α-tubulin antibody and DAPI staining. As Fig. 3, *top* shows, mitotic cells treated with the SUR A55-D70 peptide but not with the control reagent, showed several spindle abnormalities. These results are summarized in Fig. 3, *bottom*. Here, it can be seen that normal spindles were reduced by more than 50% in the mitotic cells treated with the Survivin peptide, as compared to the control. Also, this time I could see the *mini spindles* reported before by others (16, 17, 19), when interfering with the Survivin protein’s function. The *mini spindle* phenotype was 6 fold higher in the Survivin peptide-treated cells in comparison with the scrambled peptide-treated ones (28% vs. 5%). Also, but to a lesser extent, I could see other abnormal mitotic phenotypes with the SUR A55-D70 peptide, including microtubule-depleted, aberrant or multipolar spindles.

**FIG. 3.**
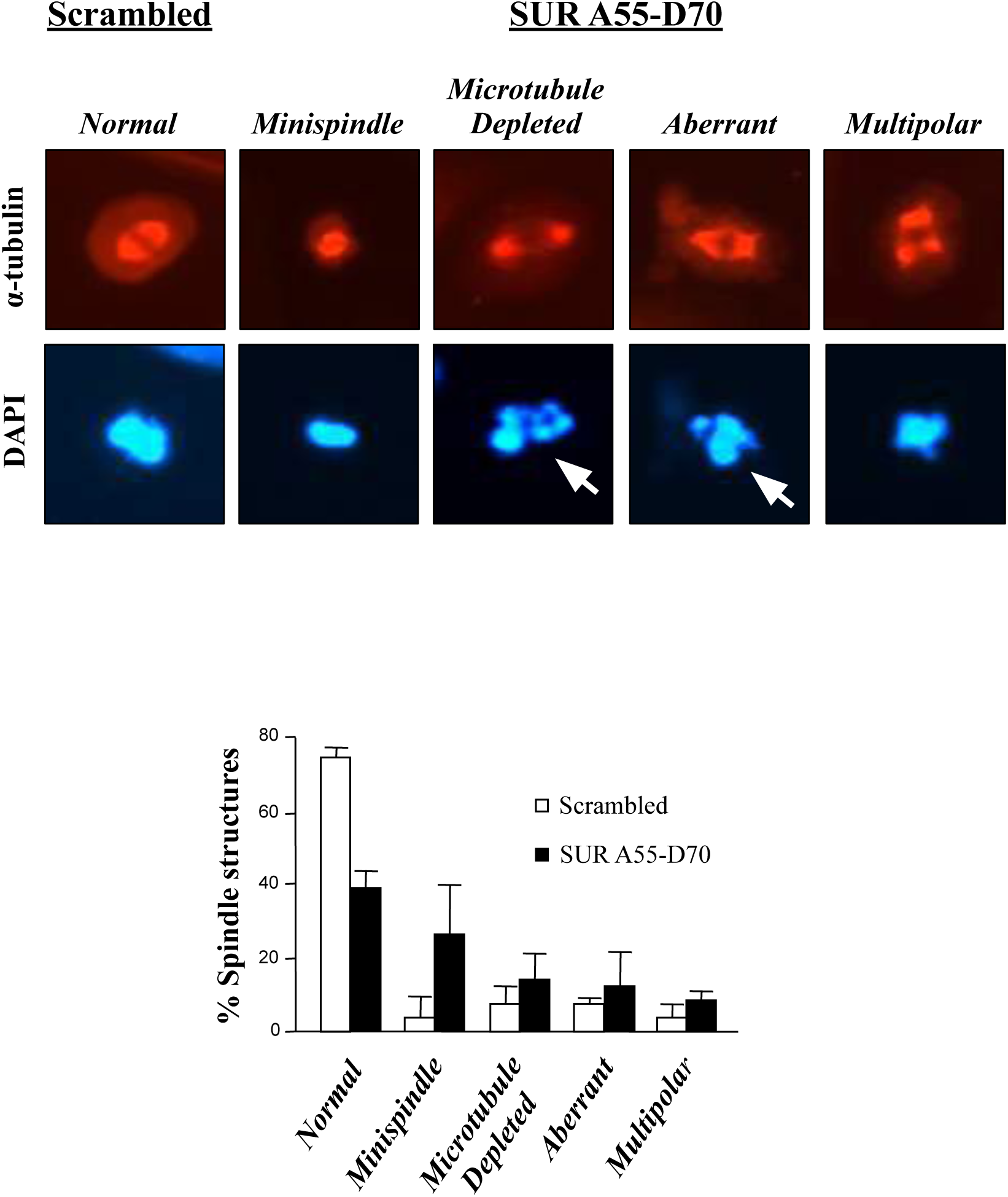
Treatment of HeLa cells with the SUR A55-D70 peptide causes spindle abnormalities. Spindle abnormalities caused by the SUR A55-D70 peptide. Asynchronous HeLa cell cultures were transfected with 50 μM scrambled or SUR A55-D70 peptide for 6 h, and cells were stained with an antibody to α-Tubulin. DNA was stained with DAPI (*top*). Mitotic phenotypes were quantified (*down*). Scrambled: 85 cells/10 fields; SUR A55-D70: 106 cells/12 fields (n=2).

When looking in detail at the DAPI staining of HeLa cells treated with the SUR A55-D70 peptide, some chromosomes appeared to mis-segregate or be fragmented (Fig. 3, *top*, white arrows). This phenotype seemed to signal cell death, and I could not observe it in the Survivin siRNA-treated samples. To investigate the possibility of the Survivin peptide causing cell death, I treated HeLa cells with the scrambled or SUR A55-D70 peptide as before, and checked their DNA content by FACS analysis after 24 h. Fig. 4A, top graphs shows that when HeLa cells were treated with 50 μM SUR A55-D70 peptide, a sub-G1 peak (9%) corresponding to cells with fragmented DNA (*<2N*) could be observed, which was almost unnoticeable in the control. To see whether I could enhance this cell death phenotype by increasing the concentration of the Survivin peptide, I incubated HeLa cells with higher concentrations of the Survivin reagent, and measured again the amount of DNA by FACS analysis. Fig. 4A, bottom graphs show that the highest peptide concentration I could reach without causing too much toxicity in the control samples was 200 μM. Under these conditions, I could observe 22% of the Survivin peptide-treated cells having fragmented DNA (*<2N*) versus 3% in the control.

**FIG. 4.**
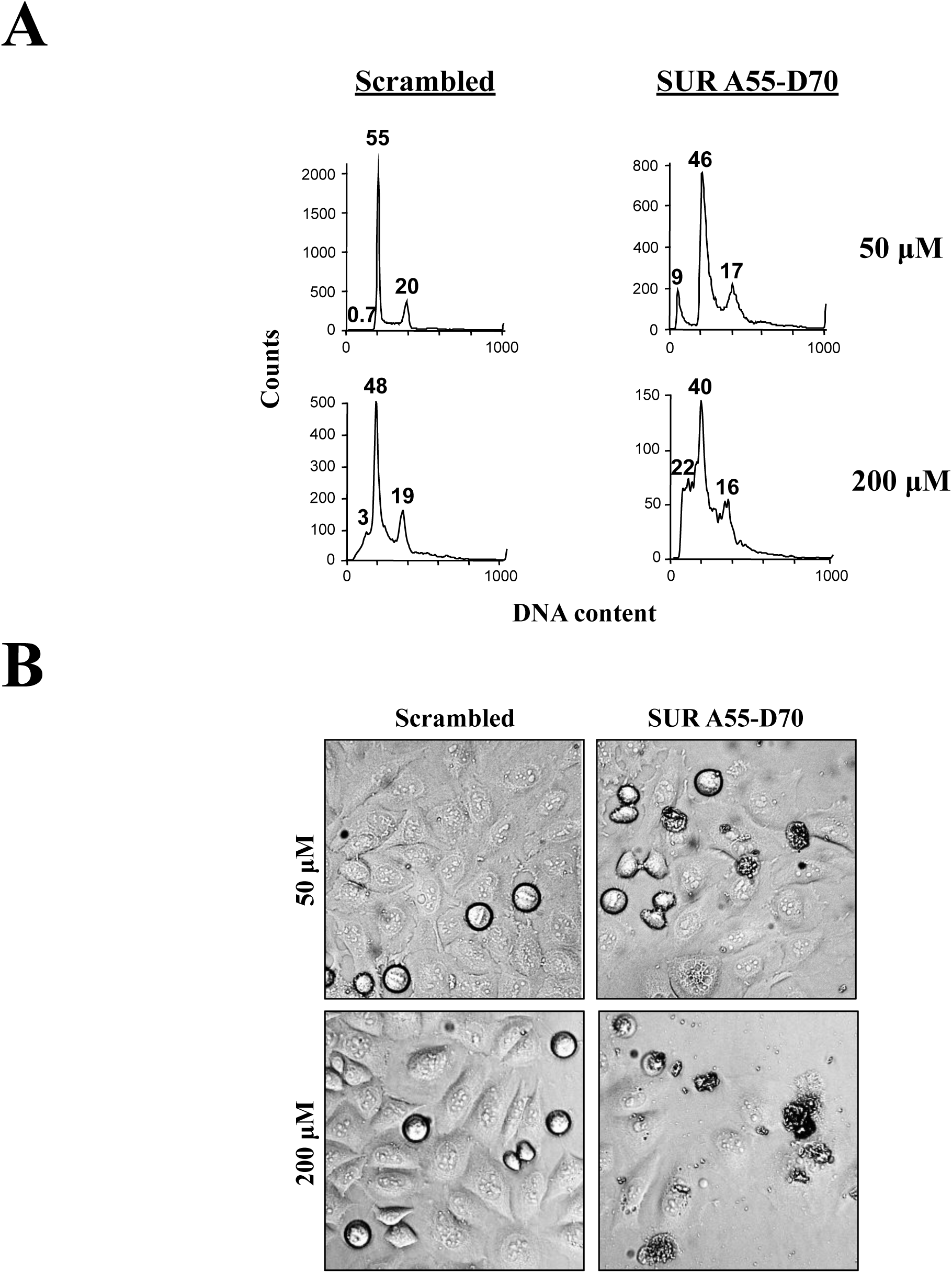
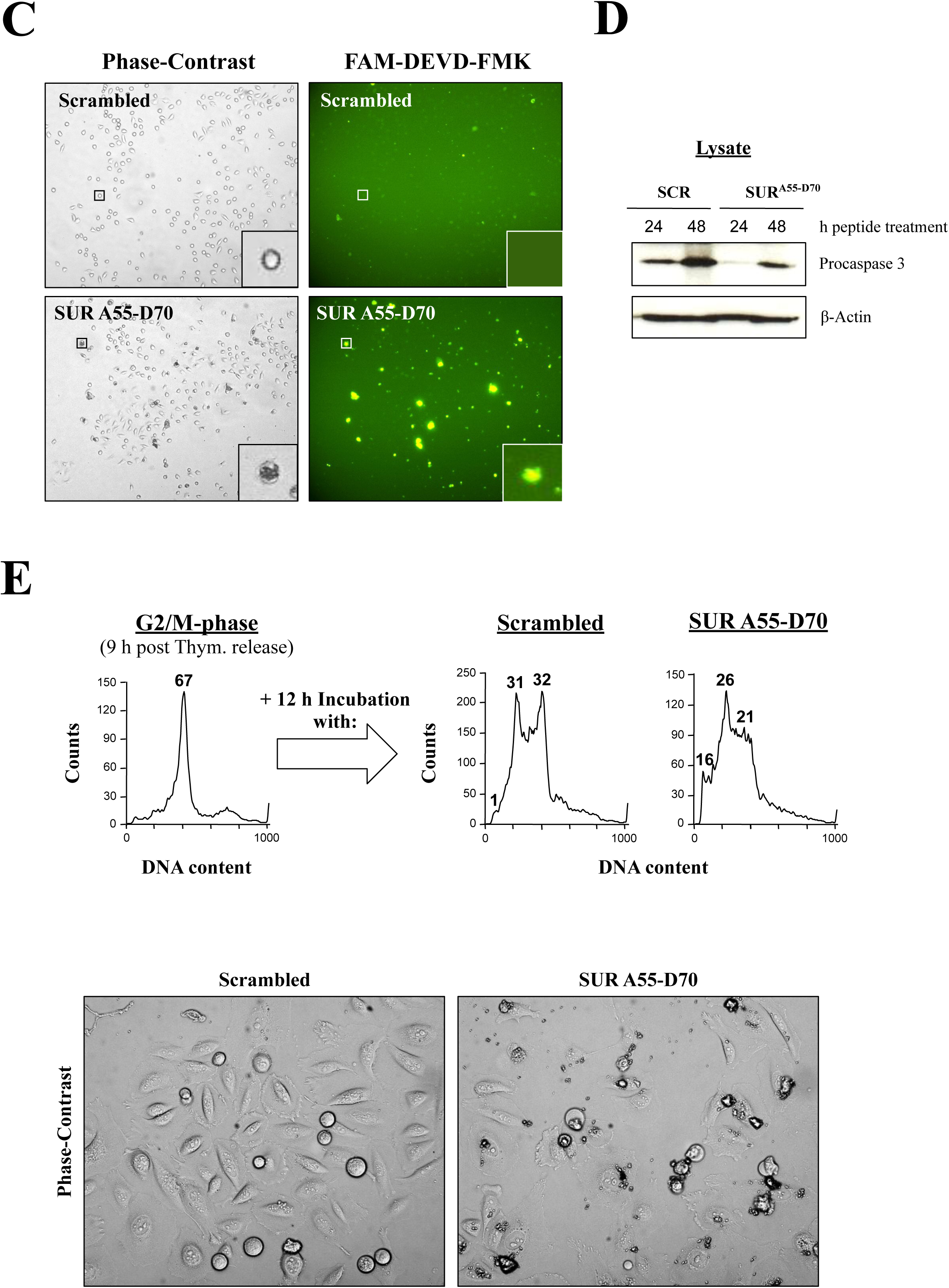
Prolonged incubation of HeLa cells with the SUR A55-D70 peptide causes apoptosis. *A*, *B*, SUR A55-D70-induced cell death. Asynchronous HeLa cell cultures were treated with 50 or 200 μM scrambled or SUR A55-D70 peptide for 24 h, and cells were analyzed by FACS (*A*) or phase-contrast microscopy (*B*). Images represent one of several experiments (n=4). The percentage of apoptotic (<*2N*), G1 (*2N*) and G2/M-phase (*4N*) cells is indicated in *A*. *C*, *D*, SUR A55-D70-induced apoptosis. Asynchronous HeLa cell cultures were transfected with 200 μM scrambled or SUR A55-D70 peptide, and cells were analyzed by phase-contrast microscopy and FAM-DEVD-FMK fluorescence microscopy (*C*) after 24h, or with an antibody to the the caspase 3 proform (*D*) after 24-48 h. Images represent one of several experiments (n=3). *E*, SUR A55-D70-induced apoptosis in synchronous HeLa cultures. G2/M-phase synchronized HeLa cells were transfected with 200 μM scrambled or SUR A55-D70 peptide for 12 h, and analyzed by FACS (*top*) or phase-contrast microscopy (*bottom*). The percentage of apoptotic (<*2N)*, G1 (*2N*) and G2/M-phase (*4N*) cells is indicated.

The sub-G1 cell population detected by FACS analysis should be easily visible under the microscope. To check this possibility, I treated HeLa cell cultures with the control or Survivin peptide at a concentration 50 or 200 μM for 24 h, and analyzed the cells by phase-contrast microscopy. As it can be seen in Fig. 4B, HeLa cells treated with the Survivin peptide but not the control one, showed signs of cell death in a dose-dependent manner, which manifested as bubbles containing a dark condensed material, or cells that looked like they had bursted and spilled out their content. Also, at the highest Survivin peptide concentration, I could see less cells under the microscope, indicating that this reagent might have cytostatic or cell lysis properties.

In order to determine the type of cell death observed in the HeLa cells transfected with the Survivin peptide, I decided to treat HeLa cell cultures similarly as above, and assay them for caspase activity. Here, two methods were used to visualize apoptosis following the peptide treatment. In the first method, cells were treated with the fluorescence probe FAM-DEVD-FMK, which labels the active caspase 3 and 7 enzymes in living cells (30). Alternatively, cell lysates were incubated with an antibody against procaspase 3, which cleavage indicates activation of the enzyme by caspases 8 and 9 (31). Fig. 4C, left panels show phase contrast images of asynchronous HeLa cell cultures treated either with 200 μM scrambled or SUR A55-D70 peptide. As it can be seen, the same cell phenotype as in Fig. 4B, right panels (i.e. cells accumulating some dense black material in their cytosol) was detected in the Survivin peptide-treated cells but not in the control-treated ones. Also, when these images were compared to the FAM-DEVD-FMK labeling (Fig. 4C, right panels), the same cells that accumulated the cytosolic dark material in the phase-contrast images, showed activation of caspases 3 and 7, as indicated by their green fluorescence. In contrast, green fluorescence could hardly be seen in the scrambled peptide-treated samples (Fig. 4C, top right panel). Identically-treated HeLa cell cultures as in Fig. 4C were used to analyze apoptosis using the procaspase 3 antibody. As Fig. 4D indicates, following the incubation of HeLa cells with the Survivin peptide for 24 or 48 h, a strong activation of caspase 3 was seen in these samples versus the control, as demonstrated by the disappearance of the caspase’s proform.

In all the above peptide experiments, the Survivin reagent targeted the Survivin-Cdk1 interface. Since this interaction mainly occurs at mitosis, I reasoned that if I enriched this cell population by using synchronous HeLa cell cultures that were allowed to progress to the G2/M-phase, and then added the Survivin peptide, apoptosis might be enhanced. To check this possibility, a synchronous HeLa cell culture released for 9 h into fresh media was prepared (G2/M-phase) (Fig. 4E, *top,* left panel), which was then incubated with either the control or SUR A55-D70 peptide for 12 h, and analyzed by FACS analysis (Fig. 4E, *top*, right panels). As it can be seen, during the time the experiment was carried out, not all G2/M-phase cells treated with the control peptide had enough time to exit mitosis (31% G1 population), and some still remained in G2/M-phase (32%), probably as a result of the treatment. However, the G2/M-phase to G1 transition in the control samples was non-traumatic, as concluded by the almost absence of cell death in these cultures (1%). In contrast to the control, although the Survivin peptide-treated samples had a similar G1 population as the control, its sub-G1 cell number was higher (16%), and seemed to originate from the G2/M-phase stalled control cells. Here, it might be possible that these stalled cells were especially sensitive to the levels of Cdk1 activity, and that they were an easy target for a reagent that affected their kinase function.

Parallel peptide-treated, synchronous HeLa cell cultures as in Fig. 4E, *top* were analyzed by phase contrast microscopy to see whether the apoptotic phenotype observed with asynchronous cells could be exacerbated. As Fig. 4E, *bottom* shows, synchronized HeLa cells targeted with the SUR A55-D70 peptide at the G2/M-phase border died more readily than the asynchronous ones, as attested by the massive amount of apoptotic cells (i.e. bubbles full of dense, dark material) seen under these conditions, which interestingly was again higher than that observed in the FACS analysis. Again, this time, a lower number of cells could be counted in the Survivin peptide-treated samples, confirming that this reagent had a cytostatic or lytic effect.

### A Survivin Asp70Ala/Asp71Ala (SUR D70A/D71A) Double Mutant Fails to Bind Cdk1 and Causes G2/M-Phase Arrest, Mitotic Abnormalities and Cell Death

To generate a second molecular antagonist of the cellular Survivin-Cdk1 complex, with which further investigate the role of Survivin at early mitosis, mutants of the Survivin domain spanning the Cdk1-binding region (i.e. amino acids Ala55 through Asp70) were generated by Alanine Scanning Mutagenesis, and screened for binding to the recombinant His-Cdk1 protein. As a result of this approach, several mutants were identified to have a reduced binding capacity to the Cdk1 kinase. These proteins included the single mutants SUR E63A, SUR E65A, SUR W67A and SUR D71A, and the double mutant SUR D70A/D71A (Fig. S5). From the mutagenesis studies, two things could be concluded. First, all the Survivin mutants, with the exception of SUR W67A, that did not bind to the Cdk1 kinase corresponded to mutations in acidic residues in the SUR A55-D70 region. Second, the best of these Cdk1-non-binding mutants was the Survivin double mutant SUR D70A/D71A. Accordingly, the SUR D70A/D71A mutant was the one selected for further studies. Finally, another interesting thing that was hypothesized, once the Survivin structure was revisited, and considered that the Survivin protein dimerizes, is that the SUR D70A/D71A mutant might act as a dominant-negative protein since the residues Asp70 and Asp71 appear on the surface of the Survivin dimer, and do not contribute to the monomers’ interface (32).

Next, the SUR D70A/D71A mutant attached to a HA-tag was tested in asynchronous HeLa cell cultures, which were analyzed by fluorescence microscopy. As it can be seen in Fig. 5A, the most prominent mitotic abnormality observed here was a phenotype that resembled the cells in prophase already reported with the Survivin siRNA (Fig. 1D). In effect, cells transfected with the HA-SUR D70A/D71A double mutant had microtubules that did not form clear spindles but surrounded their dispersed chromatin, and colocalized with the HA-tagged Survivin mutant protein (cells in Fig. 5A, right panels, labeled prophase). Interestingly, this time some HA-SUR D70A/D71A-transfected HeLa cells also appeared to have mis-segregated or fragmented DNA (Fig. 5A, white arrows), as some of the cells treated with the Survivin peptide. Coincidentally with the mitotic defects observed with the Survivin double mutant, HeLa cells expressing this protein also had a low Cdk1 activity, as measured with a Histone H1 phosphorylation assay, and a loss of binding of any Survivin protein to the Cdk1 kinase (Fig. 5B).

**FIG. 5.**
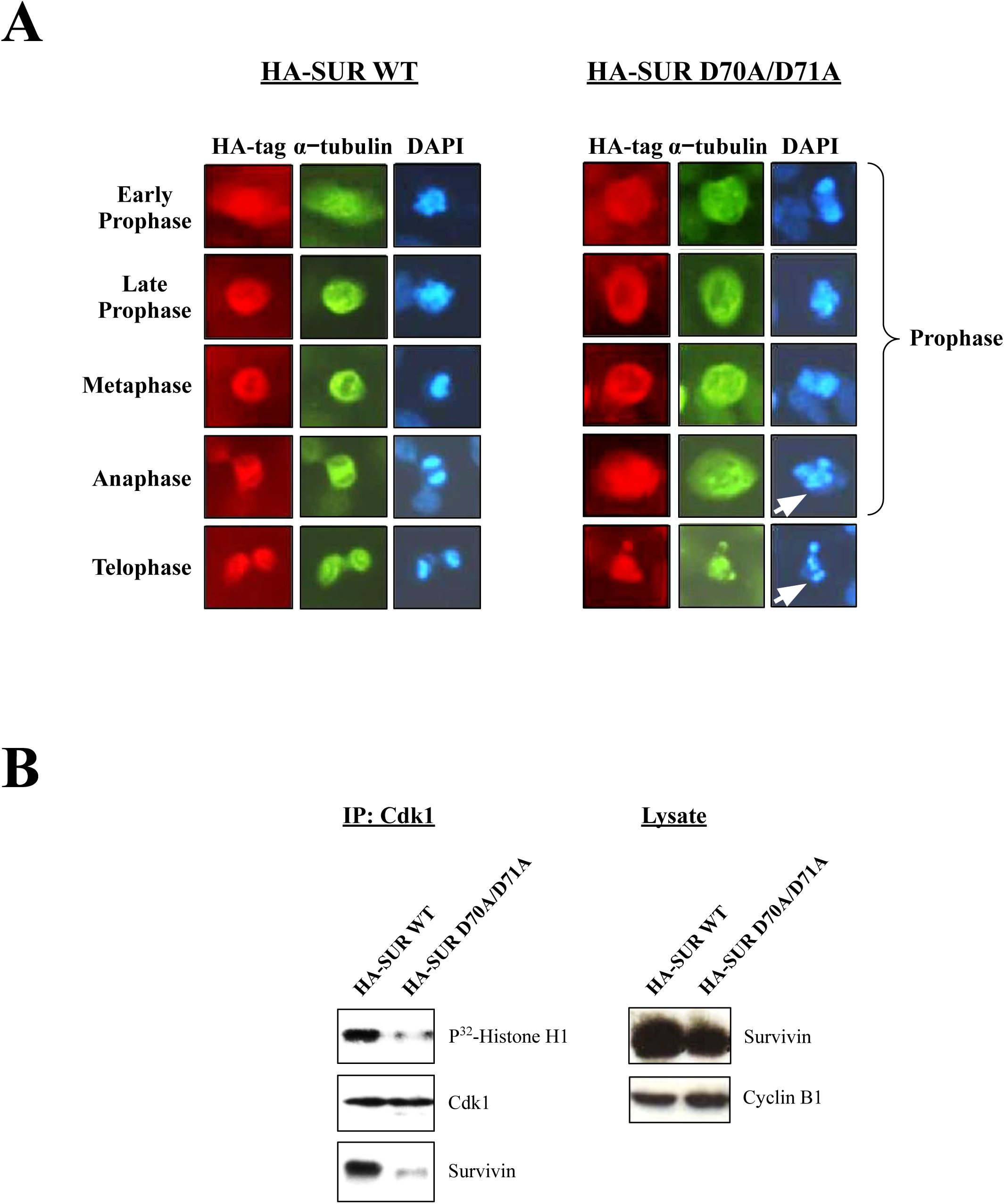
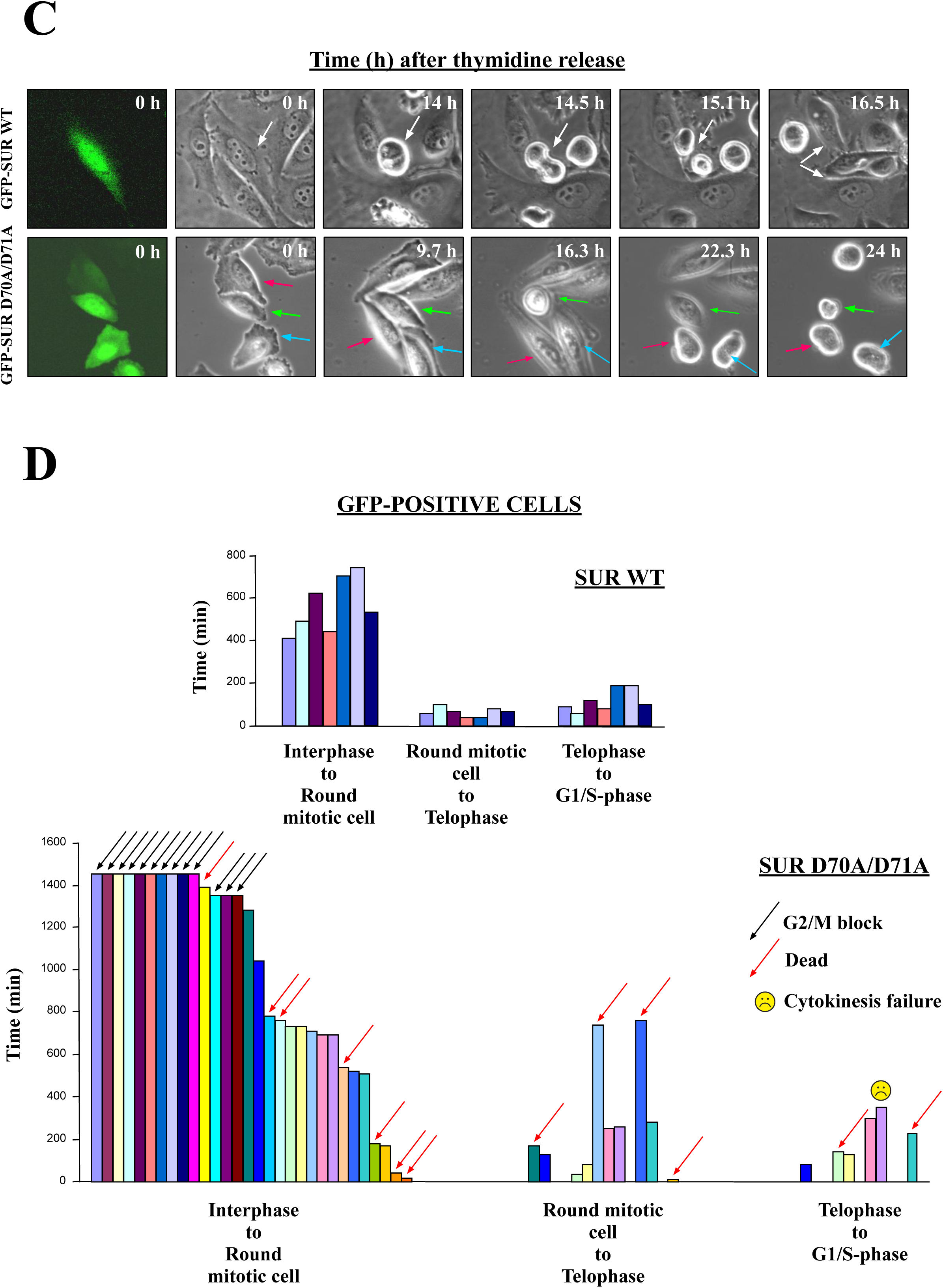
The Survivin Asp70Ala/Asp71Ala (SUR D70A/D71A) double mutant causes G2/M-phase arrest, mitotic abnormalities and cell death. *A*, SUR D70A/D71A causes mitotic abnormalities in HeLa cells. Asynchronous Hela cell cultures were transfected with constructs expressing HA-tagged wild type Survivin (HA-SUR WT) or SUR D70A/D71A (HA-SUR D70A/D71A), and subjected to fluorescence microscopy using antibodies to HA-tag or α-tubulin (*top*). DNA was stained with DAPI. *B*, Cdk1 activity in SUR D70A/D71A-expressing HeLa cells. HA-SUR WT- or HA-SUR D70A/D71A-transfected HeLa cell lysates were used to immunoprecipitate (IP) Cdk1, and pellets were analyzed in a Histone H1 phosphorylation assay (*left*), or by Western blotting as a control (*right*). *C*, *D*, Time-lapse video microscopy of SUR D70A/D71A-expressing cells. Synchronous HeLa cell cultures were transfected with GFP-SUR WT or GFP-SUR D70A/D71A. Synchronized transfected HeLa cells were released into fresh medium, and imaged every 10 min continuously for 24 h. Images were taken at the indicated time points. *C*, *bottom*, Selected cells undergoing sustained mitotic arrest (magenta and blue arrows) or cell death after mitosis re-entry (green arrow) are shown. *Top*, White arrow shows a control-transfected cell that progressed normally through mitosis. *D*, individual cells (bars), transfected with GFP-SUR WT (*top*) or GFP-SUR D70/D71A mutant (*bottom*), were quantified for time spent at each indicated mitotic transition and assigned specific phenotypes (color arrows).

In order to monitor mitotic progression in cells transfected with the Survivin double mutant, as done before for the Survivin siRNA (see Fig. 2A), I synchronized HeLa cell cultures, followed by transfecting them with either the GFP-tagged wild type Survivin protein (GFP-SUR WT) or the SUR D70A/D71A mutant (GFP-SUR D70A/D71A) in the last 12 h of their thymidine blockage, and then released them into fresh media, so they could progress through mitosis with a fully expressed transfected protein. Here, two said experiments were carried out from which 7 control and 30 Survivin double mutant-transfected HeLa cells (green cells) were monitored by time-lapse video microscopy (Figs. 5C and D). As shown in these figures, synchronized control HeLa cells (Fig. 5C, top panels and Fig. 5D, top graph) rounded around 9.5 h, which corresponded to the G2/M-phase, as previously demonstrated by FACS analysis (see Fig. 2A, *bottom*, VIII graphs). These control cells remained rounded for an average of 74 minutes before initiating telophase/cytokinesis, a stage at which they stayed for an additional 121 minutes, as average, before spreading and entering G1. Following a very different behavior (Fig. 5D, bottom graph), 13 out of the initial 30 GFP-SUR D70A/D71A-expressing HeLa cells remained flat (43%) for the 24 h that the time courses lasted. Of the other 17 green cells, 7 (23%) died early after their release into fresh media (3 cells in between 40-180 min), or later in the blockage (4 cells in between 540-1400 min or 9-24 h). The remaining 10 green cells, or 33% of the initial monitored population, rounded and entered mitosis with an average of 11.8 h or a 24% time delay in comparison with their control counterparts (average time 569 min). During mitosis, 4 out of the 10 cells that managed to escape the G2/M-phase blockage died before reaching telophase and cytokinesis (40%) (2 of them collapsing after 750 min or 12.5 h), and the other 6 progressed into the final mitotic stage, where they started splitting into two (half of these 6 cells after more than 250 min being round). Finally, during telophase/cytokinesis, 2 of the 6 cells died (33%), 1 did not complete cytokinesis, and only 3 managed to successfully complete mitosis, or 10% of the initial 30 cells used in the 2 time-lapse experiments.

A detailed microscopic analysis of individual HeLa cells expressing the Survivin double mutant showed some worth-noting behaviors. Representing these phenotypes, 3 cells labeled with a magenta, green or blue arrow are shown in Fig. 5A, bottom panels. Here, the cell labeled with the green arrow rounded very late (16.3 h), in agreement with the Fig. 5D, bottom graph, and then flattened up before dying after 24 h. To explain this behavior, I could only think that either this cell exited mitosis, and then died after trying to return to mitosis from G1, or alternatively, that the cell returned to G2, after prematurely entering mitosis, and then died when trying to re-enter prophase. Since, according to Fig. 5D, bottom graph, very few cells expressing the Survivin double mutant reached G1, I found the second explanation more plausible. Also, in support of this theory, it is worth mentioning that a return of mitotic cells to G2 has previously been reported in prophase HeLa cells that were treated with Cdk1 inhibitors (8, 14). Regarding the cells in Fig. 5C, bottom panels labeled with the magenta and blue arrows, they also rounded very late (22 h), and remained in this fashion for the rest of the time course. More interestingly, these cells had an elongated shape, similar to that seen in the RNAi experiments (Fig. 1D, bottom panels), which would be in agreement with deregulation of the Cdc25-Cdk1 axis (33).

### Survivin is Required for Recruitment and Activity of Cdk1 at the Centrosome

The results obtained so far pointed at a role of Survivin in the early activation of Cdk1. Activation of this kinase first occurs at the centrosome (6, 7), and interestingly, initial experiments with crude centrosomal preparations from synchronized HeLa cells, showed that when Survivin was immunoprecipitated (IP) from these samples, a fast-migrating Cdk1 band was bound (Fig. 6A) that has been previously ascribed to the active kinase (27). The active centrosomal Cdk1 isoform in complex with the Survivin protein first accumulated at 7 h, and preceded the FACS time point at which HeLa cells started to come out of mitosis (8 h) (Fig. S2A, *right*). Furthermore, the Survivin-Cdk1 complex continued to be visible until 11 h, or right before all the synchronized HeLa cells reached G1 (12 h) (Fig. S2A, *right*).

**FIG. 6.**
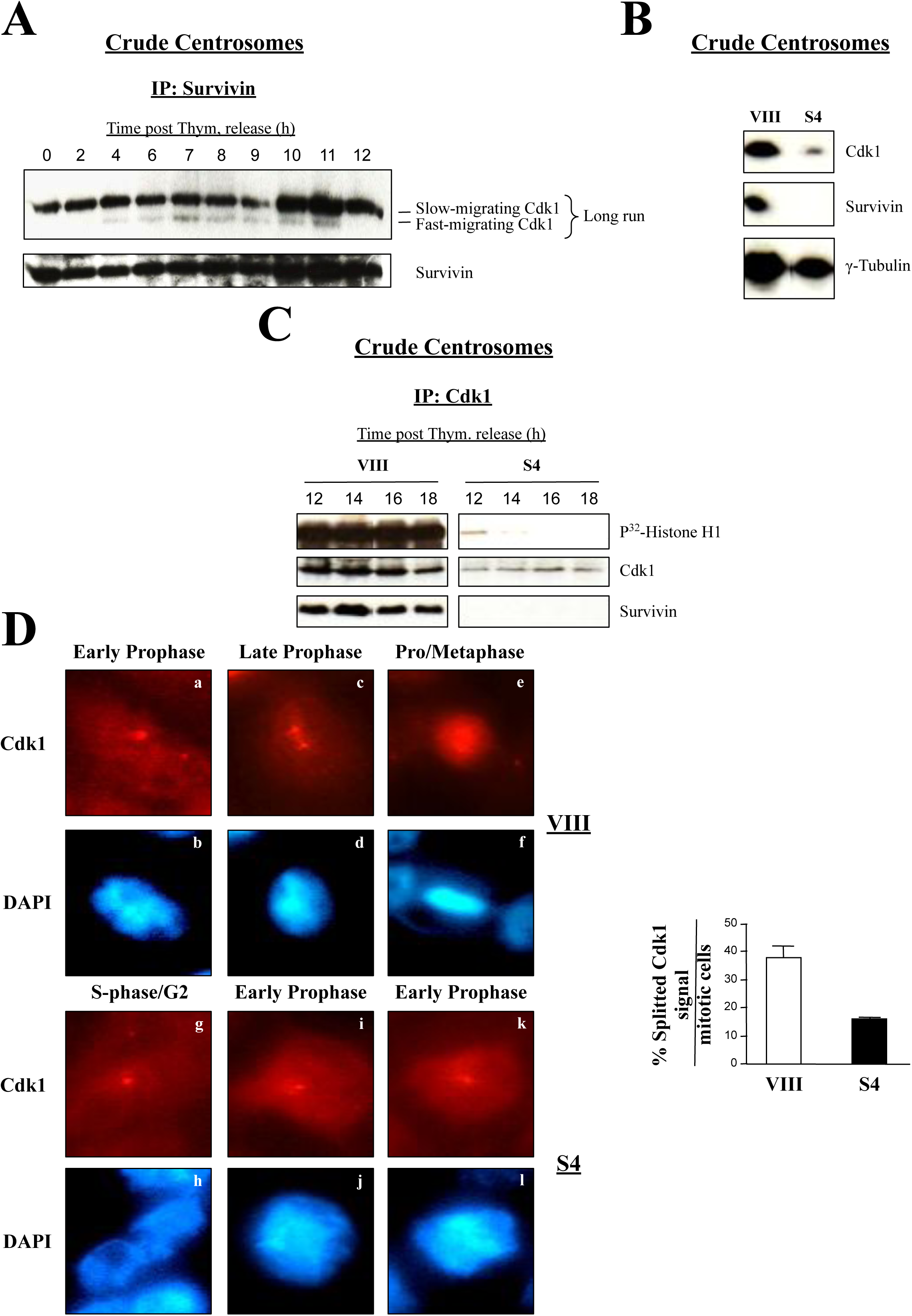
Survivin is needed to recruit Cdk1 to centrosomes. *A*, Active centrosomal Survivin-Cdk1 complex at mitosis. Centrosomes were isolated from lysates of synchronous HeLa cell cultures, subjected to Cdk1 immunoprecipitation (IP), and analyzed by Western blotting. *B*, Centrosomal Cdk1 levels in siRNA-transfected HeLa cells. Centrosomes were isolated from asynchronous HeLa cells transfected with control (VIII) or Survivin (S4) siRNA, and analyzed by Western blotting. *C*, Centrosomal Cdk1 levels and kinase activity in siRNA-treated synchronous HeLa cultures. Centrosomal preparations from synchronized HeLa cells transfected with the indicated siRNA that were collected after release into fresh media at the mentioned times, were immunoprecipitated (IP) with an antibody to Cdk1, and pellets were analyzed by Western blotting or a Histone H1 phosphorylation assay. *D*, Centrosomal Cdk1 signal in Survivin-depleted cells. siRNA-treated HeLa cells were analyzed by fluorescence microscopy using an antibody to Cdk1 (*left*). DNA was stained with DAPI. Control cells (VIII) were either in prophase (a, b and c, d) or pro/metaphase (e, f), and Survivin-depleted cells (S4) were in S-phase/G2 (g, h) or early prophase (i-l). Percentage of splitted Cdk1 signal was scored (*right*) (n=3).

In order to tackle the question of whether Survivin is required for localization and/or activation of Cdk1 at the centrosome, I treated asynchronous HeLa cell cultures with the control (VIII) or Survivin (S4) siRNA as before, and prepared centrosomes, which were used to analyze their Cdk1 content by Western blotting. As Fig. 6B shows, Survivin ablation caused a large decrease of the centrosomal Cdk1 protein, even after taking into consideration the drop in γ-tubulin. To see if this result could be reproduced in synchronous HeLa cell cultures, samples were transfected with the control (VIII) or Survivin (S4) oligonucleotide, synchronized, and aliquots were collected 12 to 18 h following cell release into fresh media, or a time that coincided with mitosis exit in the control cells (see Fig. 2A, *bottom*, upper panels). These samples were then used to prepare centrosomes from which Cdk1 was immunoprecipitated (IP), and the kinase activity was measured using a Histone H1 phosphorylation assay. Fig. 6C, left panels shows that Cdk1 was bound to Survivin in all the control samples, and that its activity was high at all times tested. In contrast, hardly any Cdk1 and no kinase activity were detected in the centrosomes of synchronized HeLa cells depleted of Survivin (Fig. 6C, right panels).

Centrosome dynamics is a well known process in eukaryotic cells (34). Briefly, in a G2 cell, the centrosome remains intact, although with duplicated centrioles. Following Cdk1 activation at prophase, the centrosome splits into two, and the daughter centrosomes gradually separate. Finally, as cells approach metaphase, the two spindle poles (i.e. daughter centrosomes) are fully separated, which coincides with the chromatin being totally condensed. Interestingly, when I started using my Cdk1 antibody in the early immunofluorescence microscopy experiments, I could identify one or two dots in mitotic HeLa cells, which seemed to mimic the above described centrosome dynamics during mitosis. This was an interesting observation, which I believed could be useful to identify the mitotic phase at which cells treated with different siRNAs were at. To this aim, I treated HeLa cells with control (VIII) or Survivin (S4) siRNA, and looked at them under the fluorescence microscope, following their labeling with the Cdk1 antibody (Fig. 6D, *left*). Here, I placed control cells transiting through mitosis in 3 categories: *a, b*, cells that contained a single Cdk1 dot and partially condensed chromatin, which I assigned to early prophase, *c, d*, cells with a partially separated Cdk1 signal, and uncompleted condensed chromatin, which I placed in late prophase, and iii) cells with two well separated Cdk1 dots, some spindle-localized Cdk1, and fully condensed chromatin, which I believed were at prometaphase or metaphase. In contrast to the control cells, a majority of the Survivin-depleted samples appeared to be in G2 (*g, h*) or early prophase (*i-l*), as indicated by their unseparated Cdk1 dot, and low chromatin condensation. To statistically determine the relevance of the unsplitted Cdk1 signal in cells treated with the Survivin siRNA (S4), I counted this phenotype from several independent RNAi experiments. As shown in Fig. 6D, *right*, lack of Survivin caused a reduction of more than 2 fold (38% vs. 15%) in the number of cells with a splitted Cdk1 signal, suggesting the implication of Survivin in centrosomal dynamics and Cdk1 activation at this organelle.

### Recombinant Survivin Induces Cdc25 Activity In Vitro

The above data pointed at a mislocalization of the centrosomal Cdk1 protein as a consequence of Survivin ablation. Since Cdk1 is first activated by Cdc25B at this location (6, 7), it was tempting to speculate that a role of Survivin might be bridging these two proteins at this organelle. If this hypothesis were true, one would expect that the lack of Survivin would cause the accumulation of the inactive Cdk1 kinase, which is phosphorylated on residues Thr 14 and Tyr 15 (PT14/PY15-Cdk1 protein) (27, 35). To check this possibility, HeLa cells were treated with the control (VIII) or Survivin (S4) siRNA, and lysates were analyzed by Western blotting using an antibody against the inactive Cdk1 form, or PT14/PY15-Cdk1 protein. As Fig. 7A, *left* shows, Survivin siRNA (S4) but not control (VIII) treated cells showed the inactive Cdk1 band. As a control, a Histone H1 phosphorylation assay of Cdk1 immunoprecipitated (IP) from the same samples was also included to confirm that the PT14/PY15-Cdk1 band correlated with low Cdk1 activity (Fig. 7A, *right*). The phosphorylated form of Cdk1 could have also been the result of Wee1 and Myt1 kinases’ activity in their role as negative regulators of cell cycle progression (5). However, Wee1 is a nuclear protein (36) and Myt1 is a membrane-bound kinase (37), so I discarded a role of these kinases at this stage.

**FIG. 7.**
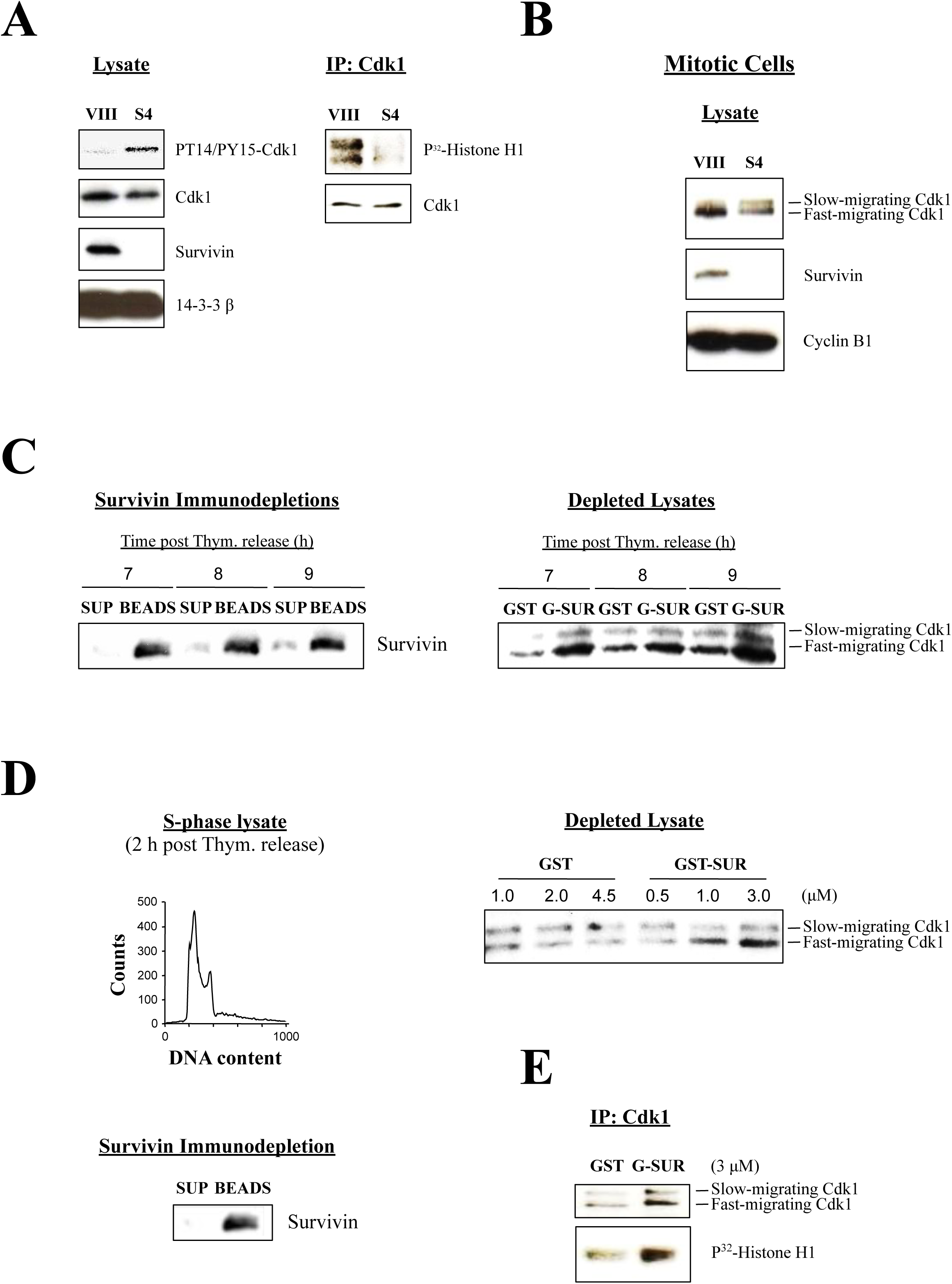
Cdc25 activity is induced by recombinant Survivin *in vitro.* *A*, Inactive Cdk1 isoform in Survivin-depleted HeLa cells. HeLa cells transfected with control (VIII) or Survivin (S4) siRNA were analyzed by Western blotting (*left*) or a Histone H1 phosphorylation assay (*right*). *B*, Impaired Cdc25 activity in Survivin-knocked down mitotic cells. Synchronized mitotic HeLa cells transfected with the indicated siRNA were collected, and analyzed by Western blotting. *C*, Induction of Cdc25 activity by recombinant Survivin *in vitro*. G2/M-phase HeLa cell lysates were depleted of Survivin (*left*), supplemented with an ATP-regenerating system, incubated with GST or GST-Survivin, and analyzed by Western blotting (*right*). *D*, *E*, Cdk1 activation by recombinant Survivin in interphase. *D*, Interphase HeLa cell lysates (*top left*), depleted of Survivin (*bottom left*), and supplemented with an ATP-regenerating system, were incubated with different concentrations of GST or GST-Survivin, and analyzed by Western blotting (*top right*), or a Histone H1 phosphorylation assay (*E*).

As already introduced in the centrosomal studies, another way to differentiate between the inactive and active Cdk1 forms is by running proteins longer during SDS-PAGE (27, 35). Indeed, it has been known for a while that the two Cdk1 isoforms appear as a doublet, an upper inactive band, which corresponds to the PT14/PY15-Cdk1 protein, and a fast-migrating active form, when these two proteins are given enough time to separate, or when the amount of polyacrylamide is increased in the gel used for SDS-PAGE. To see if I could resolve the two Cdk1 isoforms in Survivin-depleted HeLa cultures, I used control (VIII) or Survivin (S4) siRNA-treated, synchronized HeLa cells that were released for 9 h, and were therefore at G2/M-phase (see Fig. 2A), and lysates were prepared from these samples that were subjected to a long SDS-PAGE run followed by Western blotting analysis using a Cdk1 antibody. As it can be seen in Fig. 7B, a strong Cdk1 band was detected in the control cells. In contrast, in the Survivin siRNA-treated cells (S4), I could see a doublet by SDS-PAGE, consisting of a faster migrating band, similar to the one in the control sample but fainter, and also a slow-running Cdk1 form that should correspond to the phosphorylated inactive kinase (27, 35). From these and the results obtained with the PT14/PY15-Cdk1 antibody, I concluded that the lack of Cdk1 activity in the absence of Survivin seemed to be due to interference with the Cdc25-Cdk1 axis.

The above data could be more strongly supported if they could be replicated *in vitro*. To this aim, lysates from synchronized HeLa cells entering mitosis (7-9 h post thymidine release) were prepared, which were used to deplete Survivin by incubating them with a Survivin antibody that was captured with protein A beads. The Survivin immunodepleted supernatants (Fig. 7C, *left*) were then incubated with either recombinant GST or GST-Survivin protein in the presence of an ATP-regenerating system to see whether Survivin could induce the accumulation of the active Cdk1 form. As Fig. 7C, *right* shows, when Survivin-depleted HeLa cell lysates were incubated with GST, a small amount of the fast-migrating Cdk1 band could be seen that slightly increased from 7 to 9 h. In contrast, Survivin-depleted HeLa lysates that were supplemented with GST-Survivin clearly showed a higher amount of the fast-migrating Cdk1 protein at all times versus the control, which also significantly increased throughout the time course.

Next, I performed a titration experiment to see whether the accumulation of the fast-migrating Cdk1 band was dose dependent. This time, I used S-phase extracts, which should have almost no active Cdk1, and therefore provide a cleaner background. Here, it was also interesting to test whether Survivin could induce the activation of Cdk1 in interphase. This approach however had a caveat, which was that, under these conditions, Cdk1 might not be activated at all due to low upstream positive signaling in the interphase samples. To check these possibilities, Survivin-depleted HeLa cell lysates prepared from synchronized cultures collected 2 h post thymidine release (Fig. 7D, *left*) were incubated with increasing amounts of GST-Survivin or GST, and accumulation of the fast-migrating Cdk1 band was monitored. As seen in Fig. 7D, *right* increasing amounts of GST-Survivin but not GST induced the increase of the fast-migrating Cdk1 band in a dose-dependent manner. Finally, to see whether this S-phase, fast-migrating Cdk1 isoform was active, a Histone H1 phosphorylation assay was carried out. As expected, Fig. 7E shows that, even in interphase, where Cdk1 and Cdc25 activities should be low, recombinant Survivin could mount some activation of the mitotic kinase.

### Survivin is Involved in the Activation of the Cdc25B Phosphatase

As mentioned above, an explanation for the Survivin-mediated activation of Cdk1 in early prophase might be its mediation in the formation of a complex between this kinase and its activator Cdc25B. Earlier in this paper, the direct interaction between Survivin and Cdk1 (22) was confirmed (Fig. S2B), so next, I wanted to know whether Survivin could also bind to Cdc25B *in vitro*. Fig. 8A shows that when GST-Survivin was incubated with recombinant His-Cdc25B, these two proteins bound to each other. From this result, I was confident that I would find an *in vivo* Cdc25B-Cdk1 complex in the cells expressing Survivin. To my surprise, however, when I immunoprecipitated (IP) Cdk1-Cyclin B1 from siRNA-transfected HeLa cells using a Cyclin B1 antibody, I could not see Cdc25B attached to this complex in the control samples but counteractively, only in the cells that lacked Survivin (Fig. 8B). The Cdc25B-Cdk1-Cyclin B1 complex correlated with a low intensity, fast-migrating Cdk1 band, or active kinase, and the slow-moving Cdk1 form previously ascribed to the inactive kinase (27, 35). Also, this complex had to be cytosolic, as almost no Cdk1 protein was found in the centrosomes of the Survivin-depleted HeLa cells (Figs. 6B and C). Interestingly, the difference in the amount of Cdc25B bound to Cdk1-Cyclin B1 in the Survivin siRNA-treated samples replicated the amount of the phosphatase in the total lysates (Fig. 8C).

**FIG. 8.**
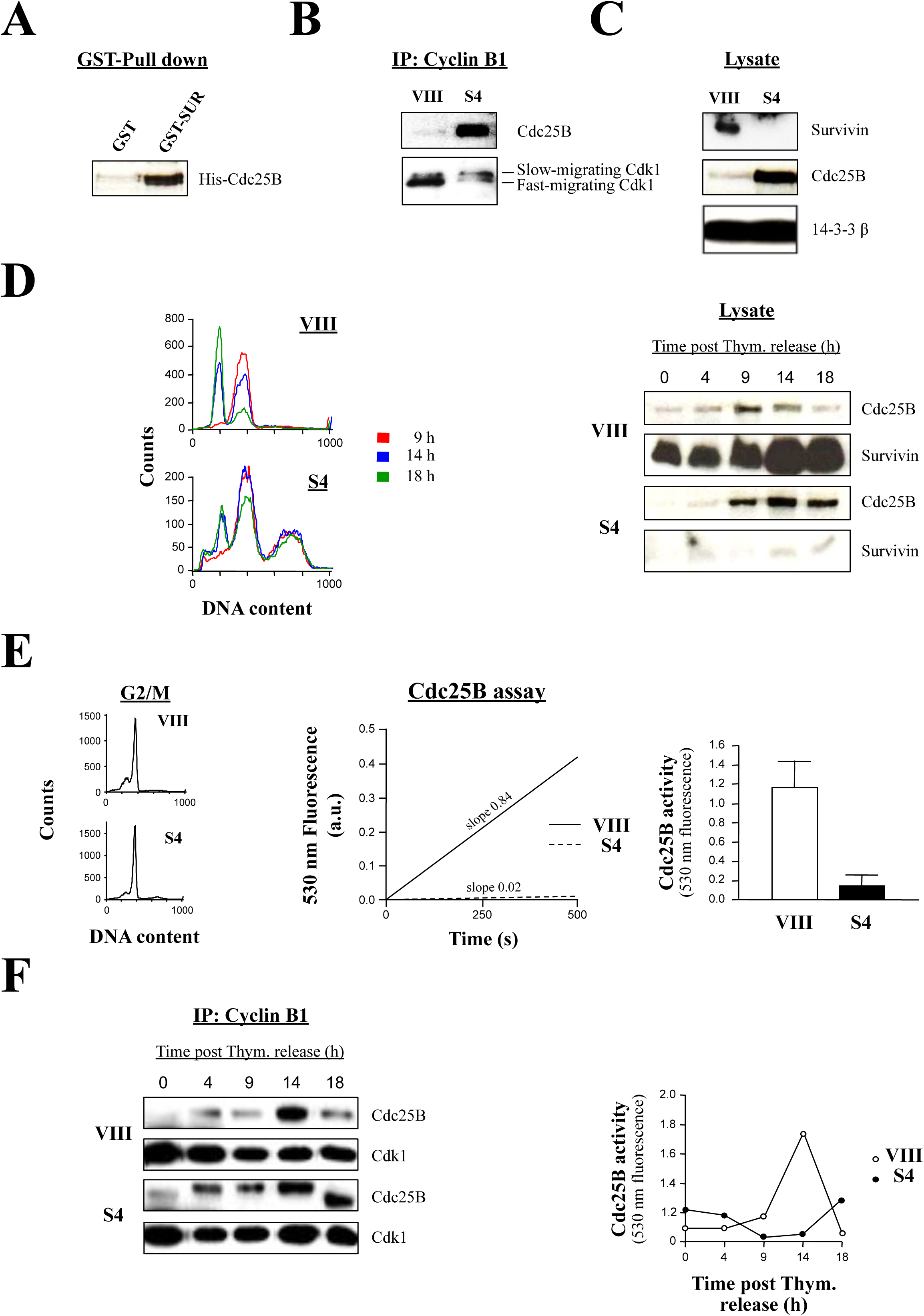
Survivin regulation of Cdc25B phosphatase activity. *A*, Survivin binds directly to Cdc25B. GST or GST-Survivin was mixed with His-Cdc25B, and analyzed by Western blotting. *B*, Accumulation of a cytosolic Cdc25B-Cdk1-Cyclin B1 complex in the absence of Survivin. Asynchronized HeLa cells transfected with control (VIII) or Survivin (S4) siRNA were collected, lysates were used to immunoprecipitate (IP) Cyclin B1, and pellets were analyzed by Western blotting. *C*, Cdc25B accumulation in asynchronous Survivin-depleted HeLa cell cultures. HeLa cells treated with the indicated siRNA were analyzed by Western blotting. *D*, Cdc25B accumulation in synchronous HeLa cultures depleted of Survivin. siRNA-treated synchronous HeLa cell cultures were analyzed by FACS (*left*) or Western blotting (*right*). *E*, Cdc25B activity at mitosis onset in siRNA-treated HeLa cells. Synchronized HeLa cells transfected with the indicated siRNA were collected at G2/M-phase (*left*), and used to immunoprecipitate (IP) Cdc25B. Cdc25B phosphatase activity in the immune complexes was measured by OMFP hydrolysis (*middle*). The result of several of these experiments (n=4) is shown on the *right*.

Next, I decided to study the behavior of the Cdc25B protein in synchronous cultures. To that aim, control (VIII) or Survivin (S4) siRNA-treated HeLa cells were synchronized as before, released into fresh media, and then their amount of Cdc25B protein was monitored as they progressed into mitosis (Fig. 8D). As it can be seen, the amount of Cdc25B increased as control cells approached the G2/M-phase checkpoint, and then declined as cells exited mitosis. In contrast, in the Survivin-depleted cells, Cdc25B reached similar levels as in the control at the G2/M-phase but then its amount was sustained for the remainder of the time course, replicating the accumulation of this protein in asynchronous cultures (Fig. 8C). Here, unexpectedly, the FACS analysis data (Fig. 8D, *left*) showed that some Survivin-depleted cells managed to escape the G2/M-phase blockage. However, this could be explained as a small amount of Survivin being expressed after 14-18 h (Fig. 8D, *right*).

The Cdc25B-Cdk1-Cyclin B1 complex observed in the asynchronous HeLa cells appeared to be inactive (Fig. 8B). A good way to confirm this result would be isolating Cdc25B from synchronous cells entering mitosis, and checking their phosphatase activity. To this aim, I immunoprecipitated (IP) Cdc25B from G2/M-phase cell cultures that had been treated with either the control (VIII) or Survivin (S4) oligonucleotide (Fig. 8E, *left*), and then the phosphatase activity was measured by employing the artificial substrate 3-O-methylfluorescein phosphate (OMFP) (38). As it can be seen in one of these experiments (Fig. 8E, *middle*), when the Cdc25B activity of Survivin-depleted cells about to enter mitosis was measured, a residual phosphatase activity was detected (0.18 a.u.), which contrasted with the strong activation of the phosphatase in the control samples (1.2 a.u.), or a 6.7 fold difference. This experiment was repeated four times and the results were very reproducible as shown in Fig. 8E, *right*.

To expand the knowledge on the Cdc25B-Cdk1 complex dynamics, and the activation of the Cdc25B phosphatase throughout the cell cycle, siRNA-treated samples were prepared as before, and their Cdk1-Cyclin B1 complexes were immunoprecipitated (IP) by using a Cyclin B1 antibody (Fig. 8F, *left*). Alternatively, Cdc25B was immunoprecipitated and used to measure its activity by OMFP hydrolysis (Fig. 8F, *right*). As Fig. 8F, *left* shows, Cdc25B formed a transient complex with Cdk1 and Cyclin B1 in control HeLa cells, which correlated with maximal phosphatase activity (14 h) (Fig. 8F, *right*), and mitosis transitioning (Fig. 8D, *left*). In contrast, in the absence of Survivin, Cdc25B remained bound to Cdk1 and Cyclin B1 from 4 to 18 h (Fig. 8F, *left*), and the activity of the phosphatase remained negligible during all time points tested (Fig. 8F, *right*).

### Only a Dominant-Positive Cdc25B Mutant Can Bypass the Early Prophase Blockage Caused by Survivin Abrogation

To confirm the effect of Survivin abrogation on Cdc25B activity, the potent capacity of the phosphatase Cdc25B to induce mitotic entry when transfected into HeLa cultures (39) could be tested in overexpression studies. Here, if Survivin had a role in Cdc25B activation, as indicated above, overexpression of the phosphatase in Survivin-depleted cells would not induce entry into mitosis. To this end, the FACS profiles of control (VIII) and Survivin (S4) siRNA-treated, synchronized HeLa cells transfected with Cdc25B were analyzed (Figs. 9A). Fig. 9A, *right*, upper graphs show that Cdc25B overexpression in control siRNA-transfected HeLa cells (VIII) resulted in an increase in the number of cells that progressed through mitosis (28% vs. 19%, or a 47% increase), as indicated by the larger G1 peak in comparison with the control (see arrows). In contrast, Survivin-ablated, synchronized HeLa cells transfected with Cdc25B remained unaltered at G2/M-phase (Fig. 9A, *right*, lower graphs), as indicated by their almost unchanged G1 peak (16% vs. 15%, or 6.7% difference).

**FIG. 9.**
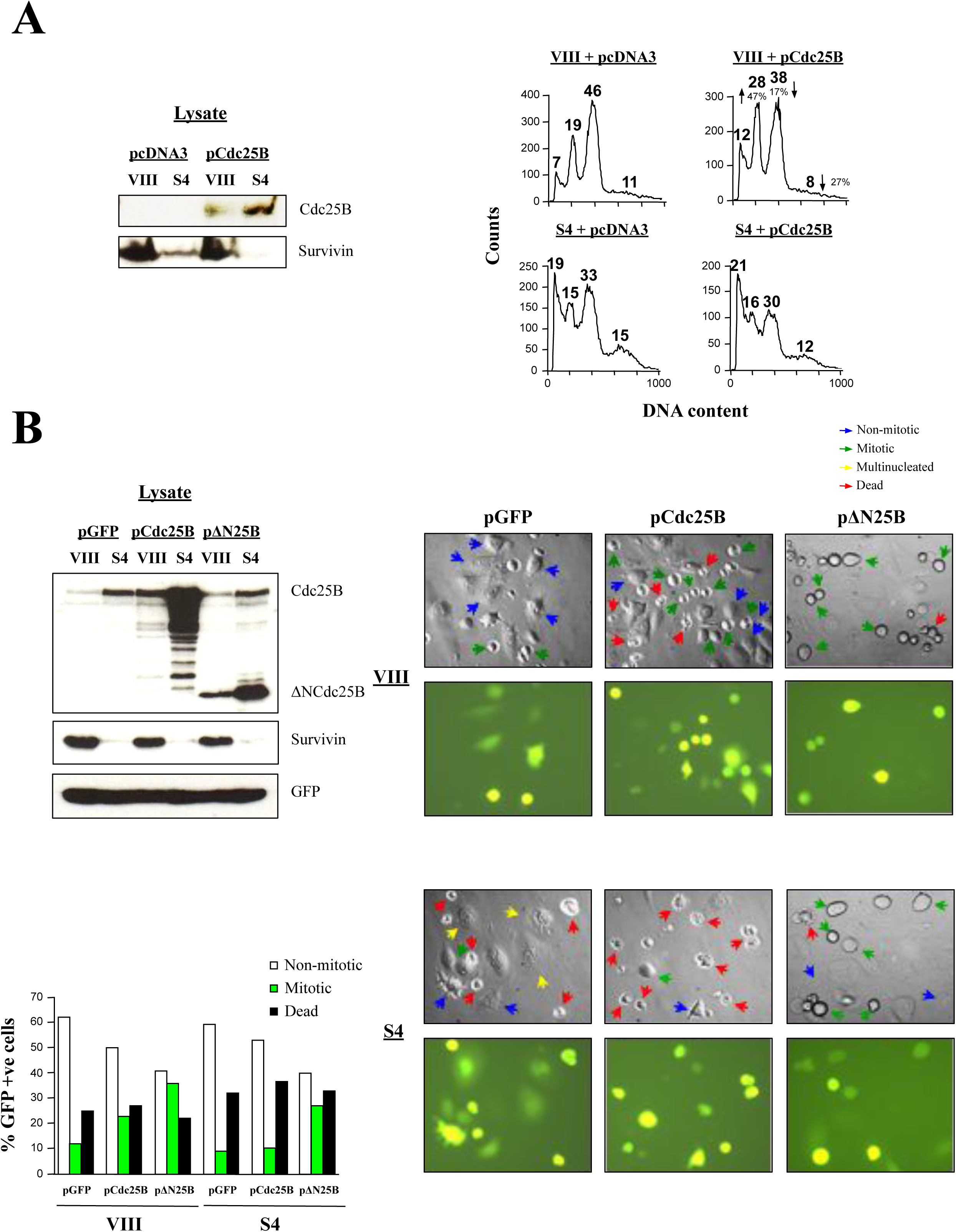
A gain-of-function Cdc25B mutant can override the blockage induced by Survivin abrogation. *A*, Cdc25B-mediated mitotic entry. HeLa cells transfected with control (VIII) or Survivin (S4) siRNA were synchronized, and then transfected with a pcDNA3 or pCdc25B plasmid, released and collected at 16 h. Cells were analyzed by Western blotting (*left*) and FACS (*right*). The percentage of cells with apoptotic (<*2N*), G1 (*2N*), G2/M-phase (*4N*) and polyploid (*>4N*) cells is indicated. Arrows and percentages indicate changes in cell populations. *B*, Gain-of-function ΔNCdc25B mutant overrides blockage induced by Survivin abrogation. siRNA-treated synchronized HeLa cells were transfected with a pGFP (control), pCdc25B (wild type Cdc25B) or pΔNCdc25B (constitutively active Cdc25B mutant) plasmid (the last 2 were also transfected with the GFP monitoring plasmid), released into fresh medium, and analyzed by Western blotting (*left*), or fluorescence and phase-contrast microscopy (*right*). Experiment was repeated 5 times with similar results. Phenotypes for one single experiment are quantified in *bottom left*.

Because Survivin appeared to be indispensable to activate the Cdc25B phosphatase, and subsequently Cdk1, I predicted that a constitutively active Cdc25B mutant being expressed in the Survivin-ablated cells would bypass the G2/M-phase blockage. To this end, a construct encoding the truncated Cdc25B protein, containing its catalytic domain but not its N-terminal regulatory region (ΔNCdc25B) was generated as before (40), and transfected together with a GFP-monitoring plasmid in control (VIII) or Survivin (S4) siRNA-treated cells (a full-length Cdc25B construct was also used as a control), and efficient transfection was detected by fluorescence microscopy. Simultaneously, I checked that the recombinant proteins were correctly expressed. As Fig. 9B, *left* shows, lysates of siRNA-treated HeLa cells transfected with the pCdc25B or pΔNCdc25B constructs showed accumulation of these proteins as expected. Here, an interesting pattern could be noticed, which was that both the exogenously-expressed full-length and truncated Cdc25B phosphatases accumulated to a large extent in the cells lacking Survivin, and followed the behavior of the cells only expressing the endogenous phosphatase under the same conditions. Individual synchronized HeLa cells transfected with the control (VIII) or Survivin (S4) oligonucleotide, and expressing the recombinant Cdc25B proteins are shown in Figs. 9B, *right*. Here, it can be seen that control (VIII) siRNA-treated HeLa cells that were transfected with pCdc25B showed more mitotic cells than cultures, which were transfected with pGFP only, in agreement with the data in Fig. 9A, *right*. In contrast, Survivin-knocked down cells transfected with pCdc25B did not enter mitosis when compared to the control but, on the contrary, died more often. On the other hand, both HeLa cells treated with control (VIII) or Survivin (S4) siRNA, followed by transfection of the constitutively-active Cdc25B mutant, readily entered into mitosis when compared to their respective controls. One representative experiment of this sort appears quantified in Fig. 9B, *bottom left*. Here, it can be seen that ΔNcdc25B was effective in bypassing the G2/M-phase blockage associated with Survivin abrogation, and concomitant low Cdk1 activity reported in these studies.

## DISCUSSION

Up to date, the Survivin protein has only been implicated in the regulation of the spindle checkpoint and cytokinesis, both functions as part of the CPC (1). In agreement with these roles, it was firmly established that Survivin abrogation first leads to a prometaphase blockage, as a consequence of errors in the correction of faulty microtubule-kinetochore attachments (2, 3). In contrast to its role in the CPC, a function of Survivin at the centrosome has not been fully investigated yet despite the fact that the use of a dominant-negative survivin mutant or a survivin antisense led to centrosomal abnormalities and disruption of a survivin-caspase-3-p21 complex at this location, which resulted in p21 degradation (18). Before this project started, I was aware of experiments carried out with *Xenopus* egg extracts (17) and HeLa cells (16, 19), which revealed that, following Survivin abrogation, a distinctive phenotype could be observed, consisting of spindles with a very short pole-to-pole distance, or *mini spindles*. In this sense, I believed that these results could poorly be explained by the sole role of Survivin in the CPC. A more plausible explanation for the *mini spindle* phenotype seemed be a function for the Survivin protein in centrosome separation, an event for which Cdk1 is necessary (20, 21) that would place Survivin at the hub where decisions to enter mitosis are made, and would nicely tied up with the activation of a default centrosomal-centered apoptotic pathway when this protein is absent (18). This possible role of Survivin, however interesting, had never been investigated before, and therefore, I decided to make it the center of my work.

### Survivin Is Required to Activate Cdk1 in Early Prophase and for Cancer Cells to Commit to Mitosis

To start this project, I tried to replicate the experiments done by others using RNAi, in which they showed that following Survivin abrogation, a prometaphase blockage ensued (2, 3). Surprisingly, when I carried out this work, I could not see many cells blocked at prometaphase, or the *mini spindles* that I was expecting, but cells that had an intact nuclear lamina (Fig. 1D). This result contradicted the *status quo* (2, 3), and could only be explained as a failure in nuclear Cdk1 activation (4, 9, 10), a phenomenon which occurs at late prophase, and determines *mitotic commitment* (13, 14). The lamina phenotype observed when Survivin was ablated could be replicated in HeLa cells treated with the Cdk1 inhibitor purvalanol A (Fig. 1E). Also, a role of Survivin in Cdk1 activation was supported by the Histone H1 phosphorylation assays of Survivin siRNA or nocodazole-treated cells, which confirmed not only that the activity of Cdk1 is low when Survivin is absent but also that the state of this enzyme under these conditions does not resemble a typical prometaphase blockage (Fig. 1G) (28).

The early prophase blockage, and low Cdk1 activity seen in asynchronous HeLa cells devoid of Survivin could also be reproduced in synchronous HeLa cell cultures, which showed a more robust impairment in mitotic progression (Fig. 2A, *bottom*, S4 graphs), and a concomitant low Cdk1 activity (Figs. 2B and C). Here, I inferred that the mitotic cells that accumulated in the G2/M-phase peak in the absence of Survivin, had to represent either G2 or early prophase cells (see asynchronous cell immunofluorescence data above). I reached this conclusion because, if there had been a large spill of G2/M-phase cells into G1, as a result of aborted mitosis (G1 cells escaping the blockage would also have a *4N* DNA content, and could have contributed to the G2/M-phase peak), these cells, or at least a fraction, should have progressed into interphase, and contributed to generating an endoreplication peak, which was not the case, due to the fact that HeLa cells possess very low levels of the p53 tumor suppressor (41), and therefore lack an effective G1 checkpoint (42). Finally, the need of Survivin for mitotic progression was further corroborated by the partial override (this partial rescue was probably due to the still presence of the Survivin siRNA in these cells) of the G2/M-phase blockage when this protein was reintroduced into the cells from which this protein was previously removed.

The use of a Survivin double mutant that could not bind to Cdk1, inhibited the Cdk1 kinase activity, and probably acted as a dominant negative protein, provided a precise timeline of the mitotic events that occur when the Survivin’s function is impaired in cells entering mitosis, and helped placing the early prophase blockage observed in the absence of Survivin within the larger picture. In effect, my results now prove that Survivin is needed to enter mitosis, and that in its absence, cells block at G2/M-phase, a result also observed with the RNAi data. Since both these Survivin antagonists inhibit Cdk1 activation, and this enzyme’s activity is needed to initiate mitosis, it is logical to think here that the blockage induced by these two reagents might have been due to a failure of the transfected cells to mount a high enough Cdk1 activity, so they could transverse the G2/M-phase threshold. Second, in the absence of Survivin, some cells manage to bypass the checkpoint and round, however, in most cases, they delay in this phase, or alternatively die, most likely as a consequence of low Cdk1 kinase activity and apoptosis activation (see discussion below). Finally, only a minority of cells lacking Survivin manage to enter cytokinesis and successfully complete mitosis.

### Centrosomal Cdk1 Does Not Accumulate in the Absence of Survivin and this May cause a Short-Circuit between this Kinase and its Centrosomal Activator/s

Cdk1 is activated by the dual-specificity phosphatase Cdc25 (5), with isoform B being the first one that acts on Cdk1 at the centrosome during early prophase (6, 7), and isoforms A and C being subsequently active at the cytosol and nucleus in a process that amplifies the initial kinase activity (5), and commits cells to mitosis (13, 14). The data in this article now shows that when the Survivin function was compromised in HeLa cells that had not entered mitosis yet (i.e. RNAi or Survivin double mutant experiments), these cells arrested at early prophase, as judged by their intact nuclear lamina and low Cdk1 activity. From this observation, I discarded a role for the isoforms A and C of the Cdc25 phosphatase behind the data presented here. In its place, I believed that centrosomal Cdc25B malfunction might have a responsibility in the lack of Cdk1 activity of the cells lacking Survivin. In agreement with this idea, when I checked the levels of centrosomal Cdk1 in Survivin-depleted cells, these were very low, and correlated with no kinase activity (Fig. 6B and C). Also, the immunofluorescence studies showed a decrease in the amount of Cdk1 at the centrosomes of Survivin-depleted cells, and a reduction in the number of cells with separated spindle poles, a phenotype previously ascribed to Cdk1 impairment (20, 21). From these results, I concluded that a failure in Cdk1 localization to the centrosome in the absence of Survivin was probably causing a short-circuit in the activation of Cdk1 by Cdc25B.

### Survivin Activates the Cdk1 Kinase Via Phosphatase Cdc25B, the Isoform that First Acts on Cdk1 at the Centrosome and Jumps Starts Mitosis

When checking the role of Cdc25 in Cdk1 activation, I found that the low Cdk1 activity in HeLa cells devoid of Survivin correlated with the detection of a previously reported inactive Cdk1 band (PT14/PY15-Cdk1) (Fig. 7A). Also, a Cdk1 protein slowly migrating in SDS-PAGE (Fig. 7B), which was previously ascribed to the inactive kinase (27, 35), could be seen. This protein could not be the result of the negative cell cycle regulators Wee1 and Myt1 since these two kinases do not localize at the centrosome but to the nucleus and endoplasmic curriculum/Golgi apparatus respectively (36, 37). Interestingly, the shift between the inactive and active Cdk1 forms could be induced in a cell-free assay where Survivin was depleted from mitotic lysates, and then added back (Fig. 7C), and also reproduced in a dose-dependent manner using interphase extracts (Fig. 7D) that phosphorylated Histone H1 (Fig. 7E). This latter result might indicate an unknown function of Survivin at interphase, which could be behind the cell death observed in cells expressing the Survivin double mutant while progressing towards the G2/M-phase checkpoint (Fig. 5D, *bottom*, lower graph).

Combining the results, which showed mislocalization of the centrosomal Cdk1 protein and detection of an inactive Cdk1 protein in the HeLa cells devoid of Survivin, with the fact that the inactive Cdk1 isoform could be shifted to the active form when Survivin-depleted lysates were incubated with recombinant Survivin, I inferred that Survivin might act by bridging Cdk1 and its Cdc25B activator at the centrosome. In support of this theory, I could see that Survivin could bind both these two proteins *in vitro* (see Figs. S2B and 8A). However, when I tried to find a Cdc25B-Cdk1 complex in cell lysates, which also contained centrosomes, of HeLa cells treated with the control siRNA, I did not get the expected result, and instead, I found a Cdc25B-Cdk1-Cyclin B1 complex in the Survivin siRNA-treated samples (Fig. 8B). According to the Histone H1 phosphorylation data, the Cdk1 bound to Cdc25B found in the absence of Survivin should be inactive, and indeed, when I looked at the immunoprecipitated Cdc25B-Cdk1-Cyclin B1 complexes in detail, I could see the slow-migrating Cdk1 band previously ascribed to the inactive kinase (27, 35). This conclusion was corroborated by directly measuring the Cdc25B activity in samples depleted of Survivin by using the artificial substrate OMPF, which showed that in the absence of Survivin no Cdc25B could be detected (Figs. E and F). From the centrosome data, it was clear that the Cdc25B-Cdk1 interaction found in the lysates from HeLa cells that did not contain Survivin had to be in the cytosol, and not the centrosome. This made me think about the possibility of Survivin acting as some kind of centrosomal anchor or scaffold for the inactive Cdc25B-Cdk1 complex, which by this means could come close to its centrosomal activator/s. Aurora A phosphorylates and activates Cdc25B at the centrosome (43). Therefore, I was tempted to speculate that Survivin might be required to bring the Cdc25B-Cdk1-Cyclin B1 complexes to the centrosome, where the phosphatase, and subsequently Cdk1, could be activated.

In the absence of Survivin, Cdc25B accumulated to a great extent (Figs. 8B, C, D and F), and remained in complex with Cdk1 (Figs. 8B and F). This was very different to the previously reported short stability of this phosphatase in cycloheximide-treated hamster BHK21 and HeLa cells where Cdc25B’s half-life was less than 30 min (44). Moreover, Cdc25B normally accumulates, and its activity precedes Cdk1 activity, then both rapidly decline after G2/M-phase (44, 45, 46), a result that I could reproduce in my studies in the control cells (Fig. 8F), as a consequence of Cdk1-dependent proteasomal degradation (47). Therefore, accumulation of the inactive Cdc25B protein in the absence of Cdk1 activity due Survivin ablation would perfectly agree with the current accepted model of Cdc25B regulation.

### A Gain of Function Cdc25B Mutant but not the Full-Lenght Phosphatase, Can Overcome the Survivin-Induced Early Prophase Blockage

Overexpression of Cdc25B normally induces mitotic entry (39, 48). However, when the exogenous Cdc25B phosphatase was expressed in Survivin-depleted G2/M-phase blocked HeLa cells, this protein could not rescue the blockage (Fig. 9A, *right*, lower right graph). This result confirmed that Survivin is required to activate Cdc25B. In agreement with this theory, when a gain-of-function Cdc25B mutant was expressed in G2/M-phase cells lacking Survivin, they managed to enter mitosis (Fig. 9B), supporting a role for Survivin in signaling through the Cdc25B-Cdk1 axis.

### Early Prophase Physiognomy and Reversibility

Following the accumulation of data in this project, it was clear that the Survivin protein is necessary to assemble a functional Cdk1 complex in early prophase. This was a valid conclusion except for the fact that only a small number of rounded cells were observed following their treatment with different Survivin antagonists (Figs. 1C and 5D, *bottom*). Searching through the literature to find an explanation for this observation, I discovered that early prophase cells do not always look round, but on the contrary, many times remain spread, resembling the physiognomy of interphase or G2 cells (13, 14). The choice between these two different appearances seems to rely on the amount of Cdk1 activity in the nucleus. Accordingly, it was quite possible that many of the cells that looked like being in G2, or interphase, following interference with the Survivin function, were actually in early prophase.

Early prophase is also a reversible phase, and this reversibility can be triggered by multiple insults. Antephase is the stage to which cells resort following the activation of a G2/M-phase checkpoint branch that responds to stress insults, and is mediated by p38 (49). This checkpoint is different to the one triggered by DNA damage, which depends on ATM and/or ATR activity (50). I would speculate here that many of the cells that entered mitosis with a low Cdk1 activity, as a result of Survivin interference, might have activated their antephase rather than their DNA damage checkpoint, and as a consequence returned to G2, hoping to return to mitosis at a better time. The antephase checkpoint only works while the activity of Cdk1 is not very high, or before nucleolar breakdown (49). This would explain why cells that were treated with the Survivin peptide could not return to G2, and continued into the apoptotic program (see below).

### Survivin Activation of Cdk1 Unveils A Second Function of this Protein in CPC Regulation

Focussing on the SUR D70A/D71A-expressing cells transitioning through mid and late mitosis, it is logical to be inclined to believe that all the abnormalities caused by this mutant at these stages were due to the role of Survivin in CPC’s regulation (1). Here, however, we do not have to forget all the evidence that supports a role of Cdk1 in the metaphase to anaphase transition, and therefore, it would be logical to reason that if Survivin participates in Cdk1 activation (this work), interference with this role might somehow affect its downstream CPC function. To refresh our memory about the importance of Cdk1 activity at the metaphase/anaphase transition, I would like to mention here a few very important pieces of data. First, INCENP is phosphorylated by Cdk1, which regulates the localization and kinase activity of Aurora-B from prophase to metaphase (51). Second, Plk1 binds to Haspin in a Cdk1-dependent manner, and reducing Plk1 activity inhibits Haspin phosphorylation of Histone H3 (52). Third, Survivin binds to phosphorylated Histone H3, and this step is necessary for CPC accumulation at the centromere (53). Interestingly, in this same work, an identical mutant as the one I used in my studies, SUR D70A/D71A, showed no binding to a phosphorylated Histone H3 peptide *in vitro* (53), implicating the same two residues, Asp70 and Asp71, in both Cdk1 activation and Survivin centromeric localization, and providing an explanation for the correct timing in the onset of both Survivin’s functions. Fourth, Borealin, a Survivin-binding partner at the centromere, is phosphorylated by Cdk1, which allows it to bind to Shughosin, and subsequently phosphorylate Histone H2A (1). Combining all this information with the results presented in this paper, it is logical to conclude that some of the abnormalities observed in the HeLa cells expressing the SUR D70A/D71A mutant while transitioning from metaphase to anaphase might have been the result of Cdk1-mediated deregulation of the CPC. Here, a few of these abnormalities might have been stalling, untimely anaphase/cytokinesis onset or mitotic slippage.

### Interference with Committed Mitotic Cells Via a Cdk1-Binding Survivin Peptide Results in Apoptosis in Cells Entering Mitosis

The Survivin peptide was made with the intention to interfere with the assembly of the Survivin-Cdk1 complex. This means that this reagent should not affect interphase cells, but cells in mitosis where this complex should be abundant and the kinase active (i.e. committed mitotic cells). In this sense, treating HeLa cell lysates from nocodazole-treated HeLa cells with the Survivin peptide caused complex disassembly, as expected, and loss of Cdk1 activity (Fig. S4B), proving that Survivin needs to be bound to Cdk1 for the kinase to be active. Also, treatment of HeLa cells with the Survivin peptide correlated with several spindle abnormalities, including the *mini spindle* phenotype that inspired this work, in committed mitotic cells, as judged by their attempt at forming a DNA segregation apparatus (5). Interference with Cdk1 activity in cells that have disassembled their nuclear envelope (i.e. committed) (8), and progressed to prometaphase with a fully active Cdk1 kinase (13, 14) might lead to conflicting orders and mitotic disarray (Fig. 3), as Cdk1 is needed for spindle formation at this stage (20, 21, 54, 55). Another consequence of interfering with Cdk1 activity in committed mitotic cells might be *mitotic abortion* and cell death (15). This is precisely what happened in HeLa cells treated with the Survivin peptide. Here, I want to remind us about the fact that this phenotype was not observed in the Survivin siRNA- or Survivin double mutant-treated cells, and this might have to do with these cells having enough time to wait at the G2/M-phase checkpoint following the Survivin antagonist treatment. In support of the vulnerability of tumoral cells committed to mitosis when targeted with a Cdk1 inhibitor, cancer cells and tumors in a cancer xenograft model were very sensitive to a combination of taxol and purvalanol A but not to the reverse order of the treatment (56), indicating that when challenged cancer cells are allowed to rest at the G2/M-phase checkpoint, they can avoid cell death.

Regarding the cell death observed when HeLa cells were treated with the Survivin peptide, it has been shown that Cdk1-Cyclin B1 activity is required to phosphorylate procaspase 8 (57) and procaspase 9 (58), and subsequently block the extrinsic (57) and intrinsic (58) apoptotic pathways in cancer cell lines. Here, blockage of Cdk1 activity by Cyclin B1 RNAi or the Cdk1 inhibitor RO-3306 (57), or short-circuit of the kinase by the expression of non-phosphorylatable caspase proforms (57, 58), led to apoptosis. These were precisely the damages sustained by HeLa cells that were treated with the Survivin peptide reagent in my studies. In effect, when I used a FAM-DEVD-FMK or an antibody against procaspase 3, I could detect caspase 8 and 9 activity triggered by the Survivin peptide (see Figs. 4C and D). From these results, the Cdk1 kinase rises as some kind of mitotic protective shield but also as a factor, which might contribute to tumorigenesis if deregulated. Curiously, these have been roles traditionally assigned to Survivin, and we may now have the reason behind these phenotypes.

Even though there has been a lot of discussion regarding the two seemingly roles of Survivin in cancer regulation, namely: the one in mitosis and the one in apoptosis (for reviews on these two different views see 59 and 60), not much effort has been done trying to unify these two fields. In this respect, I propose here for the first time an integrating theory in which the two roles of Survivin can coexist. In this model, Survivin’s main function in cancer cells would be controlling mitotic entry through its regulation of Cdk1 activity. This control over Cdk1 by Survivin would have downstream repercussions on its latter role at the CPC, and ultimately would lead to apoptosis when the Cdk1 is deregulated through interference with the Survivin function at the checkpoint via biological or chemical insults.

### Survivin Binds to the αC/β4 Loop in Cdk1, a Region Involved in Protein-Protein Interactions, Binding of the Hsp90-Cdc37 Complex and Regulation of Kinases’ Activity

The Survivin peptide spanning the Cdk1-binding region (Figs. S3A and C) comprises several negatively-charged residues that reside on the Survivin dimer’s acidic surface (32). In fact, even though I used the SUR D70A/D71A mutant as a Survivin antagonist due to this protein not being able to bind Cdk1, both the GST-SUR 71-142 truncated protein, which could not bind the kinase (Fig. S2C), and the alanine point mutant SUR D71, which slightly did (Fig. S5, *bottom*), suggested that only the Asp70 is crucial for binding to Cdk1. From these observations, I predicted that Survivin should probably recognize a region in the Cdk1 kinase with a net positive charge, and indeed, I was able to narrow down the Survivin-binding sequence in the Cdk1 kinase to amino acids Lys56 through Met85, a region that includes the kinase’s αC/β4 loop, which contains several basic residues (Fig. S3B). Residues 66 through 85 were also part of the original Cdk1 sequence recognized by Survivin, however this kinase region harbors numerous hydrophobic amino acids, and therefore I found it unlikely to be part of the Survivin-binding domain.

The αC/β4 loop is a conserved structural motif present in all eukaryotic protein kinases, which bridges the αC helix in the N lobe and regions at the top of the C lobe (61). In most kinases, this loop spans 8 amino acids, which follow the consensus sequence: L-x-H-P-N-T-V-x, where x represents any amino acid (62). Functionally, it has been hypothesized that the αC-β4 loop may act as a molecular brake for protein kinases by maintaining auto-inhibitory interactions via hydrogen bonding with the hinge region in the C lobe (63, 64, 65). In support of this theory, many mutations have been found at this location, which confer constitutive kinase activity and/or drug resistance, and play a direct role in cancer progression (62). The αC-β4 loop is also a site for protein–protein interactions, and here, one of the most important roles of this loop is the recognition of the molecular chaperone Hsp90 and its co-chaperone cdc37 (66, 67), proteins that together promote the proper folding of 60% of the human kinome (67, 68). By cryoEM structure of an Hsp90-Cdc37-Cdk4 complex, a general mechanism of Hsp90-Cdc37 action has been postulated (69), where a conserved H-P-N motif in Cdc37 would mimic the turn within the kinase αC-β4 loop, and push against the kinase αE helix in the C-lobe, preventing the contact between the kinase N and C lobes. As a result of this intermolecular interaction, the kinase would be initially stabilized, and would expose other motifs to which Hsp90 could bind. From all this information, it would be tempting to speculate that Survivin might also have a role in the regulation of Cdk1 inter-lobe movement and its activation. In support of this view, Survivin recapitulates some of Cdc37’s features. In effect, apart from binding to the αC-β4 loop region as already stated, Survivin, like Cdc37, binds to the N-terminus of the Hsp90 chaperone (70, 71). Also, I could detect a complex between Survivin, Hsp90 and the active Cdk1 protein form at mitosis (Fig. S6). For these reasons, it seems reasonable to believe that coinciding with mitotic onset, Survivin might function as some kind of Hsp90 co-chaperone. Another possibility might be that Survivin binds the chaperone machinery, once the Cdk1 complex has been assembled, and brings it close to its activator/s at the centrosome. Still, a third possibility might be that Survivin unloads the Hsp90-assembled Cdk1 complexes, and delivers them to its centrosomal activator/s. In this regard, this work showed that when Survivin was absent, an inactive cytosolic Cdc25B-Cdk1-Cyclin B1 complex accumulated, and here, it would be interesting to find out whether this complex is bound to Hsp90. A function of Survivin in mediating Cdk1 through Hsp90 is however a little bit controversial, as it encounters those who claim that Cdk1 is not a Hsp90 client (68), and others that support the opposite (67, 72, 73).

### Future Strategies Regarding Cancer Treatment Arising from these Studies

Survivin has been called a *cancer gene* (74). This paper now shows that this title may have in part been earned due to an up-to-date unreported role of Survivin in the activation of the Cdc25B-Cdk1 complex. In this regard, there is plenty of research showing a correlation between Cdc25B signaling and tumorigenesis, and it would make perfect sense that cancer cells hijack this pathway in order to control mitotic entry. The connection between Cdc25B and cancer has led to an overwhelming effort trying to develop efficient Cdc25B inhibitors (75). These inhibitors however most of the time target the Cdc25B catalytic domain, and here, an exciting alternative might be finding small molecules that disrupt the Survivin-Cdc25B interaction. Here, another area worth clarifying in the future should be finding the Survivin upstream partner, which links with this protein and the Cdc25B-Cdk1 complex.

In this work, I have also shown that the absence of Survivin leads to an early prophase blockage due to low Cdk1 activity, and propose that this phenotype is very sensitive to events, which override the G2/M-phase checkpoint. Therefore, a combination of drugs that inhibit Cdk1 activity (this work), and promote at the same time mitotic entry (here the archetype might be the taxol-purvalanol A cocktail (56)), might be a very effective way to combat cancer in the future.

Allosteric inhibitors are also on the way that bind to the αC-β4 loop in Cdks (76), and this paper reinforces the importance of this site by proving that Survivin is another protein that binds to this epitope. From different studies, the αC-β4 loop is now rising as some sort of a hub through which kinases communicate with other proteins, and signals can be sent through different pathways. Therefore, elucidating the exact function of this region in kinases, and the role of the chaperone machinery at this location, may help refine the design of new drugs against cancer.

To end, I would like to reiterate the connection between Cdk1 activity and CPC’s regulation, and the possible function of Survivin at this stage. In this context, Survivin might operate as a coordinator of upstream and downstream mitotic events, which would lead to smooth progression through mitosis.

## MATERIALS AND METHODS

### Plasmids, Cloning, Recombinant Proteins and Peptides

Full-length human cDNAs encoding Cdk1 (BC014563) and Cdc25B (BC051711) were obtained from Open Biosystems. Cdk1 and Cdc25B cDNAs were subcloned in frame with the N-terminal His-tag into pRSETa (Novagen, Merck Biosciences). Cdc25B cDNA was also subcloned into pcDNA3 without (pCdc25B). pGST-Survivin (plasmid containing a Survivin sequence cloned in frame with the N-terminal GST-tag into pGEX (Pharmacia)) and pGFP-Survivin (plasmid made in pcDNA3 including an N-terminal GFP sequence (pGFP)) were available in the laboratory. Deletion mutants for His-tagged Cdk1 (residues 1-43, 1-56, 1-85, 1-183, 9-85, 36-85 and 182-297), Cdc25B (residues 391-580 (ΔNCdc25B)) and GST-Survivin (residues 1-70, 15-142, 38-142, 55-142, 71-142 and 81-142) were generated by PCR. Alanine mutagenesis of the Survivin sequence (pGST-Survivin) was carried out using the QuikChange^R^ Site-Directed Mutagenesis Kit (Stratagene). Plasmid encoding GFP-Survivin D70A/D71A (pGFP-Survivin D70A/D71A) was made by cloning the Survivin D70A/D71A sequence obtained by alanine mutagenesis into pGFP. All constructs were confirmed by DNA sequencing. His-tagged and GST-fusion proteins were expressed in BL21 *E. Coli* strain, and isolated by affinity chromatography on Ni^2+^-charged agarose (Novagen, Merck Biosciences), or glutathione agarose (Sigma-Aldrich). A peptide comprising residues Ala55 throughout Asp70 in Survivin (AQCFFCFKELEGWEPD), and a scrambled version of the same peptide (EPCWDECFAEKFQGFL), both containing a Biotin moiety and a HIV tat cell-permeable sequence at the N-terminus, were synthesized using f-BOC chemistry, purified by HPLC and analyzed by mass spectrometry (Peptron, Inc., Daejeon, South Korea or W. M. Keck Biotechnology Research Center, Yale University, School of Medicine).

### Cell Culture and Synchronization, and Transfections

Cervical carcinoma HeLa cells (ATCC, Manassas, VA) were maintained in culture as described (16). Cells were synchronized by incubation with 2 mM thymidine (G1/S-phase) or 10 μM nocodazole (M-phase). In some experiments, cells were treated with the Cdk1 inhibitor purvalanol A (10-20 μM) for 16 h. Gene targeting by small interfering RNA (siRNA) was carried out with a control or a Survivin-siRNA oligonucleotide characterized in earlier studies (16). When using synchronized cultures, cells were first treated with the appropriate siRNA, and then incubated with thymidine for 48 h. Cells were then released into fresh medium, and samples taken at increasing time intervals. Transfection of adenovirus vectors encoding GFP-Survivin (pAd-Survivin) or GFP (pAd-GFP) was performed as indicated (77). For plasmid transfection, cells (50-70% confluency) were incubated with 1-2 μg of plasmid in 1 mL Optimem-I and 4 μL lipofectamine/well for 4 h. When siRNA and pAd vectors were co-transfected, cells were first incubated with control or Survivin siRNA, and synchronized in G1/S-phase for 48 h. Cells were then released into fresh medium, transfected with the appropriate pAd vector, and harvested at increasing time intervals. In the case of transfections with siRNA and Cdc25B plasmids, cells remained in thymidine for 32 h after the siRNA treatment, plasmids were transfected in the presence of thymidine for 4 h, and cells were finally released 12 h post-transfection and harvested as indicated.

### Preparation of Cell Lysates and Isolation of Centrosomes

Cells were lysed in 1 mM HEPES, pH 7.2, 0.5% IGEPAL CA-630, 0.5 mM MgCl_2_, 0.1% β-mercaptoethanol, 50 mM NaF, 1 mM Na_3_VO_4_ plus protease inhibitors for 30 min on ice. The cell lysate was pipetted up and down five times, centrifuged at 2,500 g for 10 min, and supernatant was adjusted to 10 mM HEPES, pH 7.2 final concentration. For isolation of crude centrosomes, 1 μg/mL DNAse I was added to the lysate, and incubation proceeded for 30 min on ice. The mixture was then overlaid on a 60% sucrose cushion, containing 10 mM HEPES, pH 7.2, 0.1% Triton-X100 and 0.1% β-mercaptoethanol, and centrifuged at 10,000 g for 30 min. The interface of the sucrose cushion containing enriched centrosomes was collected for the various experiments. Protein concentration was estimated by the BCA assay (Pierce).

### GST-Pull Downs, Immunoprecipitations and Antibodies

Pull-down experiments with His-tagged proteins mixed with GST, GST-Survivin or GST-Survivin mutants were carried out as described (78). For streptavidin-binding experiments, 50 μg of biotinylated peptides were bound to 25-50 μl of a 50:50 slurry of streptavidin agarose (Sigma-Aldrich), mixed with recombinant proteins or cell lysates, and analyzed by Western blotting. Immunoprecipitation using primary antibodies (10 μg/mL), and 200-400 μg of total or fractionated cell extracts were carried out as described (78). The following antibodies to Cdk1, PT14/PY15-Cdk1, 14-3-3 β, Lamin B (all from Santa Cruz Biotechnology), Cyclin B1 (PharMingen), Survivin (NOVUS Biologicals), α-tubulin (clone B-5-1-2) (Sigma-Aldrich) and His-tag (Invitrogen) were used. Protein bands were visualized by enhanced chemiluminescence (Amersham Biosciences).

### Analysis of Cdk1 Activity

Cell lysates in 10 mM HEPES, pH 7.2, 0.5% IGEPAL CA-630, 0.5 mM MgCl_2_, 0.1% β-mercaptoethanol, 50 mM NaF, 1 mM Na_3_VO_4_ plus protease inhibitors were used to immunoprecipitate Cdk1. Kinase activity was assayed as described (22). For the *in vitro* Cdk1 activation experiments, synchronized HeLa cell lysates were prepared by incubation for 30 min on ice in 10 μM HEPES, pH 7.4, 10 μM KCl, 1.5 mM MgCl_2_ and 2 μM DTT plus protease inhibitors, and passing the cells ten times through a 27 ½ G needle. Lysates were centrifuged at 2,500 g for 10 min, and supernatants were adjusted to 10 mM HEPES, pH 7.4, 100 μM KCl and 1 mM DTT. Survivin was depleted in the lysates by rotation with 25-50 μl of a 50:50 slurry of protein A agarose (Boehringer Ingelheim) bound to Survivin antibody overnight at 4°C. Survivin-depleted supernatants were supplemented with an ATP-regenerating system (2 mM ATP, 10 mM phosphocreatine and 3.5 U creatine kinase per 100 μL lysate), and incubated with 0.5-3 μM GST or GST-Survivin for 1 h at 30°C. For the peptide interference of Cdk1 activity *in vitro*, lysates were prepared from nocodazole-treated cells as described for the Cdk1 activation experiments but this time the endogenous Survivin protein was not depleted, and samples were incubated with 1-5 μM peptides for 1 h at 30°C. All reactions were terminated by a 10-fold dilution in a buffer containing 50 mM NaF and 1 mM Na_3_VO_4_. Phosphorylated Cdk1 proteins were immunoprecipitated, resolved by SDS-PAGE and identified by Western blotting. Cdk1 activity was assayed as described (22).

### Cdc25B Activity Assay

Cdc25B protein was immunoprecipitated from cell lysates as described (78), and phosphatase activity was assayed by 3-O-methyl fluorescein phosphate (OMFP) hydrolysis at 30°C in assay buffer (100 mM Tris, pH 8.2, 40 mM NaCl, 1 mM DTT and 20% glycerol) (38). Fluorometric detection was carried out at 485 nm excitation and 530 nm emission wavelengths (Photon Technology International, Inc.). Fluorescence slopes were calculated and normalized against values obtained with control, non-binding IgG.

### FACS Analysis

DNA content in asynchronous or thymidine-synchronized HeLa cell cultures was determined by FACS analysis (propidium iodide staining and flow cytometry) as described (22). Data were quantified using the FlowJo cell cycle analysis software.

### Analysis of Apoptosis

DNA integrity was analyzed by scoring the sub-G1 population (*<2N*) in FACS analysis. Caspase activity was detected by incubating siRNA transfected HeLa cells with a fluorescein-conjugated caspase inhibitor (FAM-DEVD-FMK). Alternatively, HeLa cell lysates were analyzed by Western blotting using an antibody to procaspase 3.

### Fluorescence Microscopy

Transfected HeLa cells seeded onto optical grade glass coverslips were fixed in ice-cold 4% paraformaldehyde/PBS, pH 7.2 for 1 h, washed and permeabilized in 0.2% Triton-X100/PBS, pH 7.2 for 30 min at 22°C, except when they were transfected with GFP constructs, in which case, cells were directly analyzed under the microscope. Coverslips were blocked in 3% BSA/0.1 % Tween-20/PBS, pH 7.2 for 30 min at 22°C, and incubated with primary antibodies appropriately diluted in blocking buffer for 1 h at 22°C. Cells were incubated with Alexa Fluor (Invitrogen) secondary antibodies for 1 h at 22°C in the dark. When cells were transfected with biotinylated peptides, they were directly incubated with Texas red-streptavidin, following fixation and permeabilization. DNA was stained with 10 μg/mL DAPI (Sigma-Aldrich). Coverslips were analyzed using an inverted fluorescence microscope (Olympus IX71). Pictures were taken using a microscope-attached camera (Nikon), and data was computer-analyzed by the IPLab software.

### Time-Lapse Video Microscopy Analysis

Thymidine synchronized HeLa cells seeded onto Lab-Tek two-chambered borosilicate coverglass slides (Nalge Nunc International) were transfected with the appropriate GFP plasmid, released into fresh medium, and imaged using an inverted microscope (Zeiss) with a 20X numerical aperture 0.5 objective lens, a spinning-disk confocal scan head (Perkin Elmer Life and Analytical Sciences) and an Orca-ER cooled CCD camera (Hamamatsu) under conditions of CO_2_ exchange (0.5 L/min) at 37°C. Time-lapse imaging was conducted using a multidimensional image software (Metamorph 6.3r6) to acquire images every 10 min for 24 h, observe multiple x,y-coordinates at every time point and acquire six optical sections over a 10 μm z-series (2 μm step size) at each coordinate. A z-stack of GFP images was taken at the start and end of each time-lapse experiment.

## ACKNOWLEDGEMENTS

The author wants to thank Dr. Stephen J. Doxsey for his help in the time-lapse video microscopy studies, and his laboratory colleagues for their input. This work was supported by NIH grants CA78810, CA90917 and HL54131.

## SUPPLEMENTAL MATERIAL

### SUPPLEMENTAL FIGURE LEGENDS

**FIG. S1.**
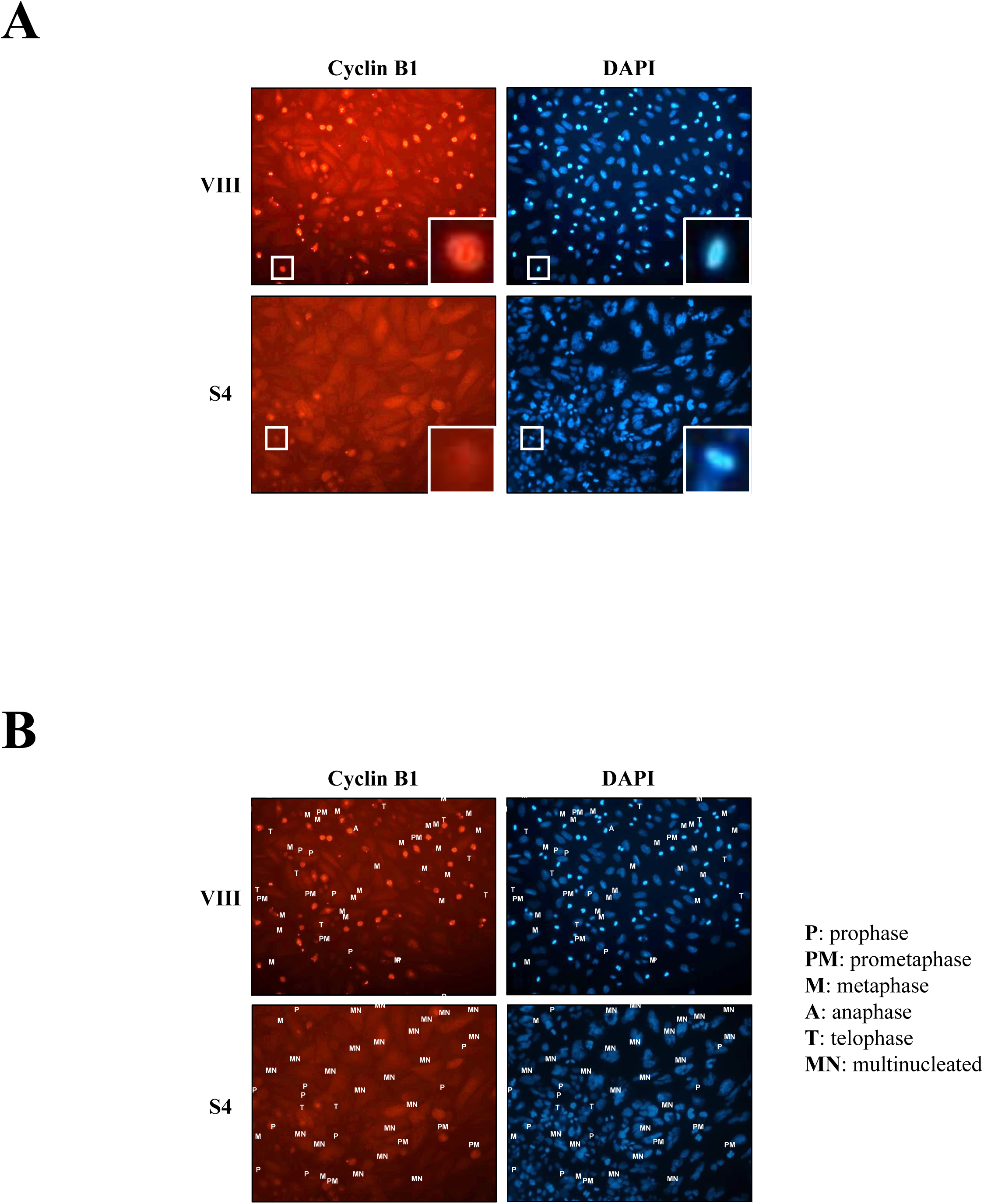
Loss of Survivin causes a reduction in mitotic cells. *A*, Cyclin B1 staining in siRNA-treated HeLa cells. HeLa cells transfected with control (VIII) or Survivin (S4) siRNA were analyzed by fluorescence microscopy with an antibody to Cyclin Β1. DNA was stained with DAPI. *B*, Cell cycle phase of Cyclin B1-positive siRNA-treated HeLa cells. Dimmed pictures in *A* were labeled according to their mitotic phase.

**FIG. S2.**
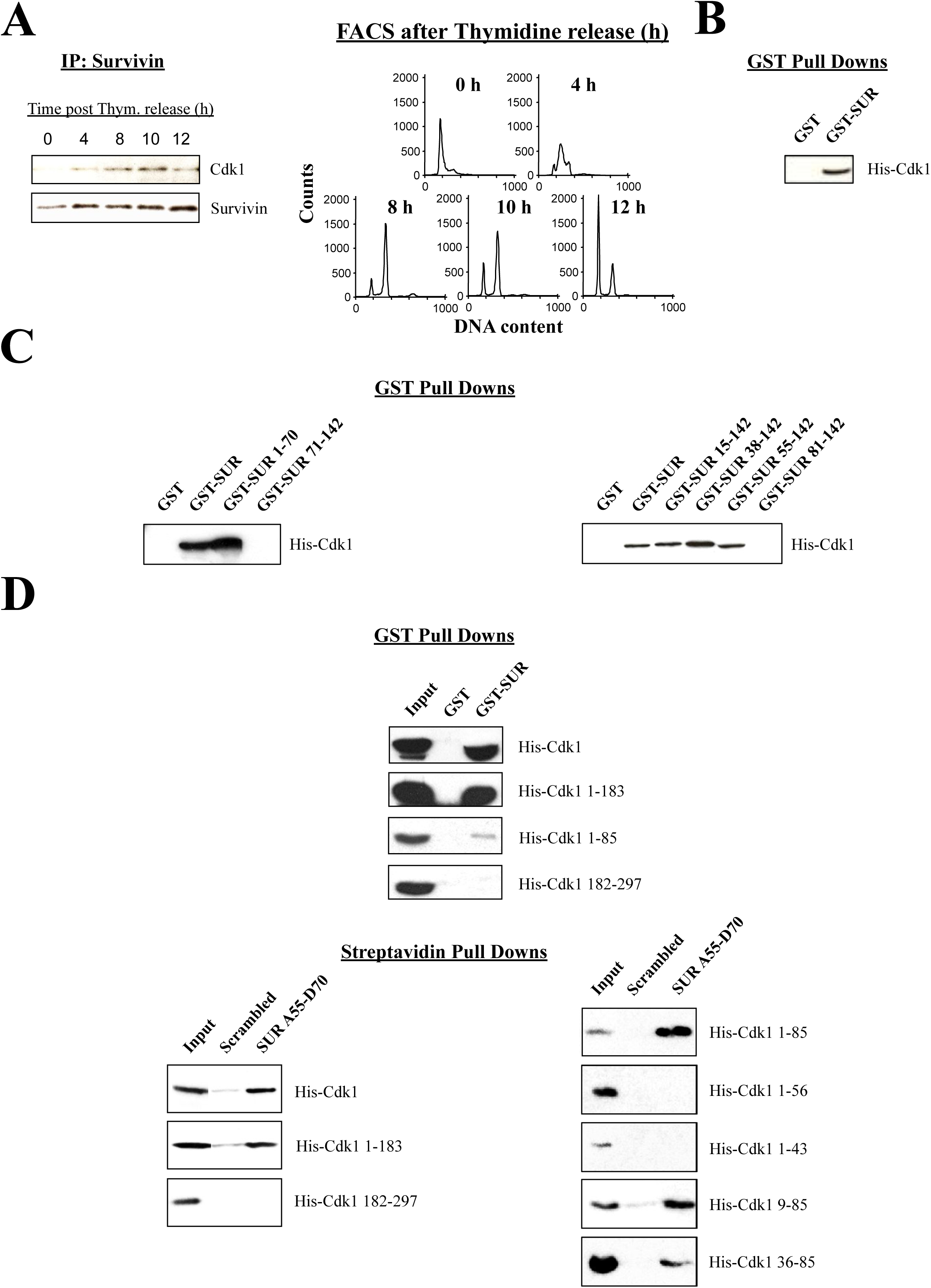
Characterization of the Survivin-Cdk1 interaction *in vitro*. *A*, Mitotic interaction of Survivin and Cdk1 *in vivo*. Synchronized HeLa cells were released into fresh medium, harvested at the indicated time intervals, immunoprecipitated (IP) with an antibody to Survivin and pellets were analyzed by Western blotting (*left*). Replica samples were analyzed by FACS (*right*). *B*, Survivin directly interacts with Cdk1 *in vitro*. Recombinant GST-Survivin and His-Cdk1 were used in a pull down experiment. *C*, Cdk1-binding Survivin region. GST or the indicated GST-Survivin deletion mutant (numbers correspond to amino acid residues in the Survivin sequence) was mixed with His-Cdk1, and bound proteins were analyzed by Western blotting. *D*, Survivin-binding Cdk1 region. GST-Survivin (*top*) or biotinylated SUR A55-D70 peptide (*bottom*) was mixed with His-tagged full-length or the indicated Cdk1 fragment followed by Western blotting.

**FIG. S3.**
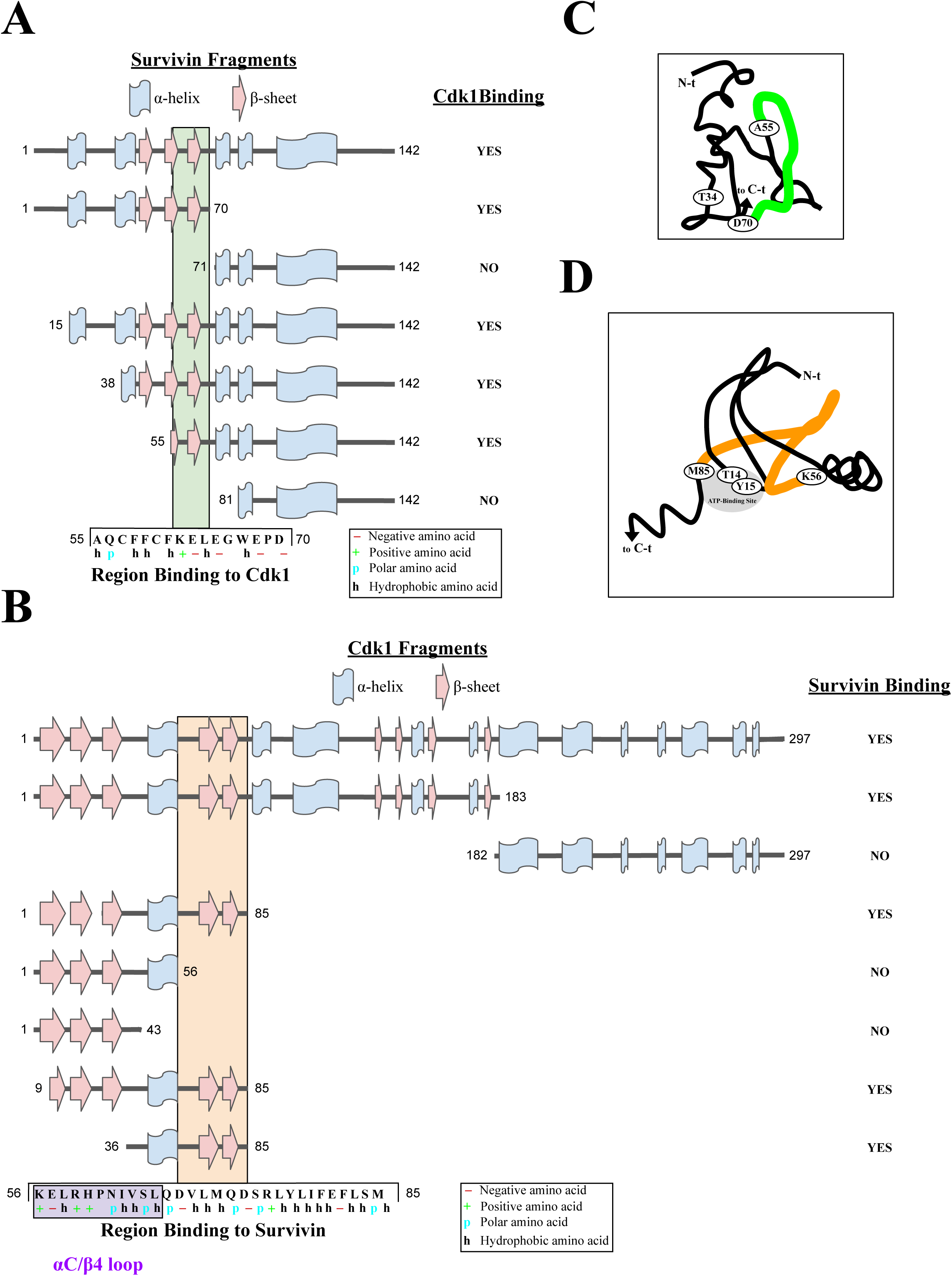
Regions involved in the Survivin-Cdk1 complex. *A*, Cdk1-binding Survivin region. *B*, Survivin-binding Cdk1 region. *C*, SUR A55-D70 peptide superimposed on the N-terminal Survivin region. *D*, Survivin-binding Cdk1 region superimposed on N-terminal Cdk1 region. All protein sequence and functional information used to generate Survivin’s and Cdk1’s graphs and cartoons was extracted from UniProt.

**FIG. S4.**
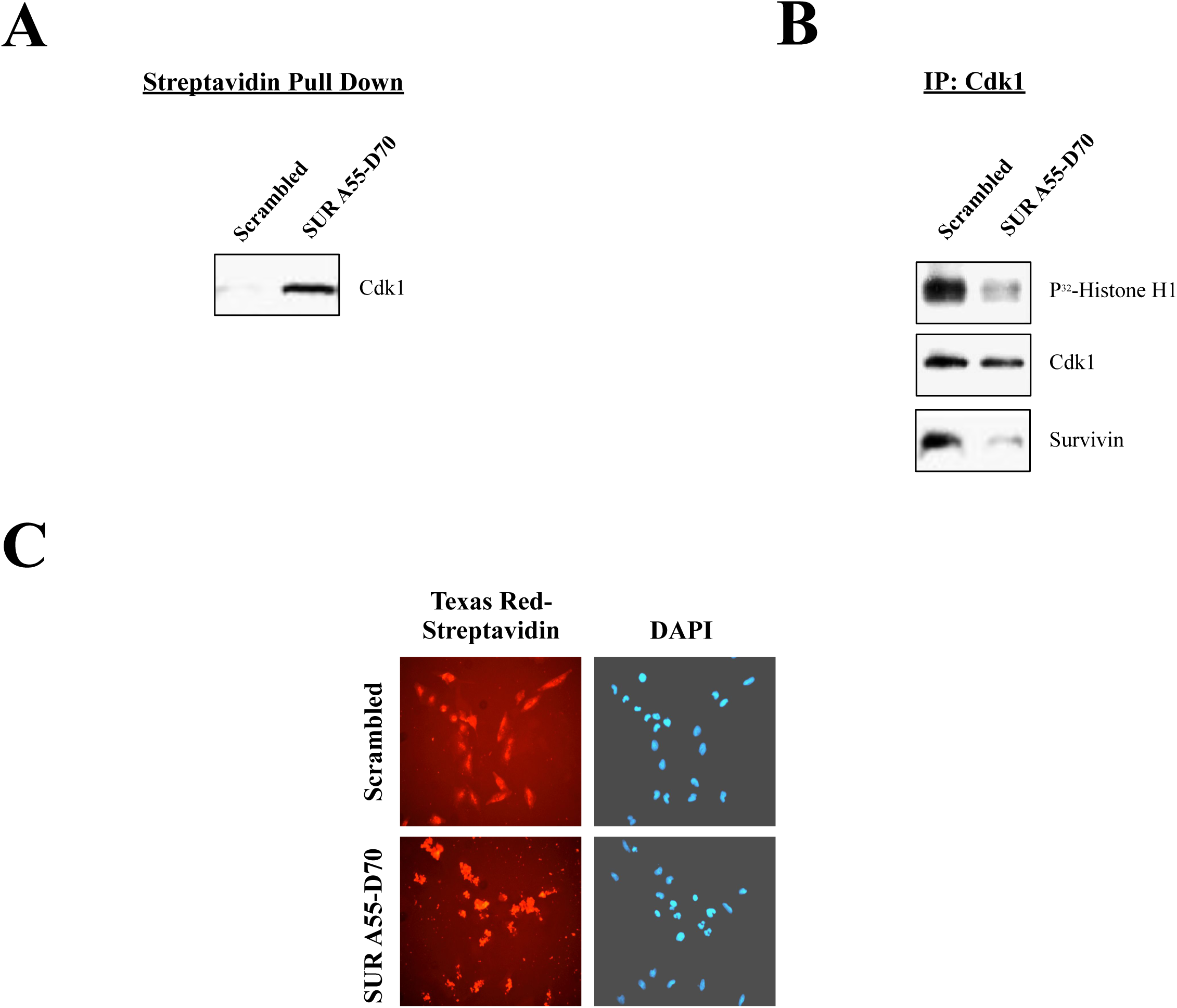
Characterization of the SUR A55-D70 peptide. *A*, SUR A55-D70 pull down of endogenous Cdk1. HeLa cell lysates were used to pull down Cdk1 using a biotin-conjugated scrambled or Survivin A55-D70 (SUR A55-D70) peptide. Protein complexes were recovered bound to streptavidin agarose and analyzed by Western blotting. *B*, SUR A55-D70-mediated disassembly and loss of kinase activity of a Survivin-Cdk1 mitotic complex. Lysates were prepared from nocodazole-treated HeLa cells, supplemented with an ATP-regenerating system, and incubated with the scrambled or SUR A55-D70 peptide (5 μM) for 1 h at 30°C. Cdk1 was immunoprecipitated (IP), and immune complexes were analyzed in a Histone H1 phosphorylation assay or by Western blotting. *C*, Peptide delivery to HeLa cell cytosol. HeLa cells were incubated with biotinylated, HIV tat-scrambled or SUR A55-D70 peptide, and analyzed by fluorescence microscopy using Texas red-streptavidin. DNA was stained with DAPI.

**FIG. S5.**
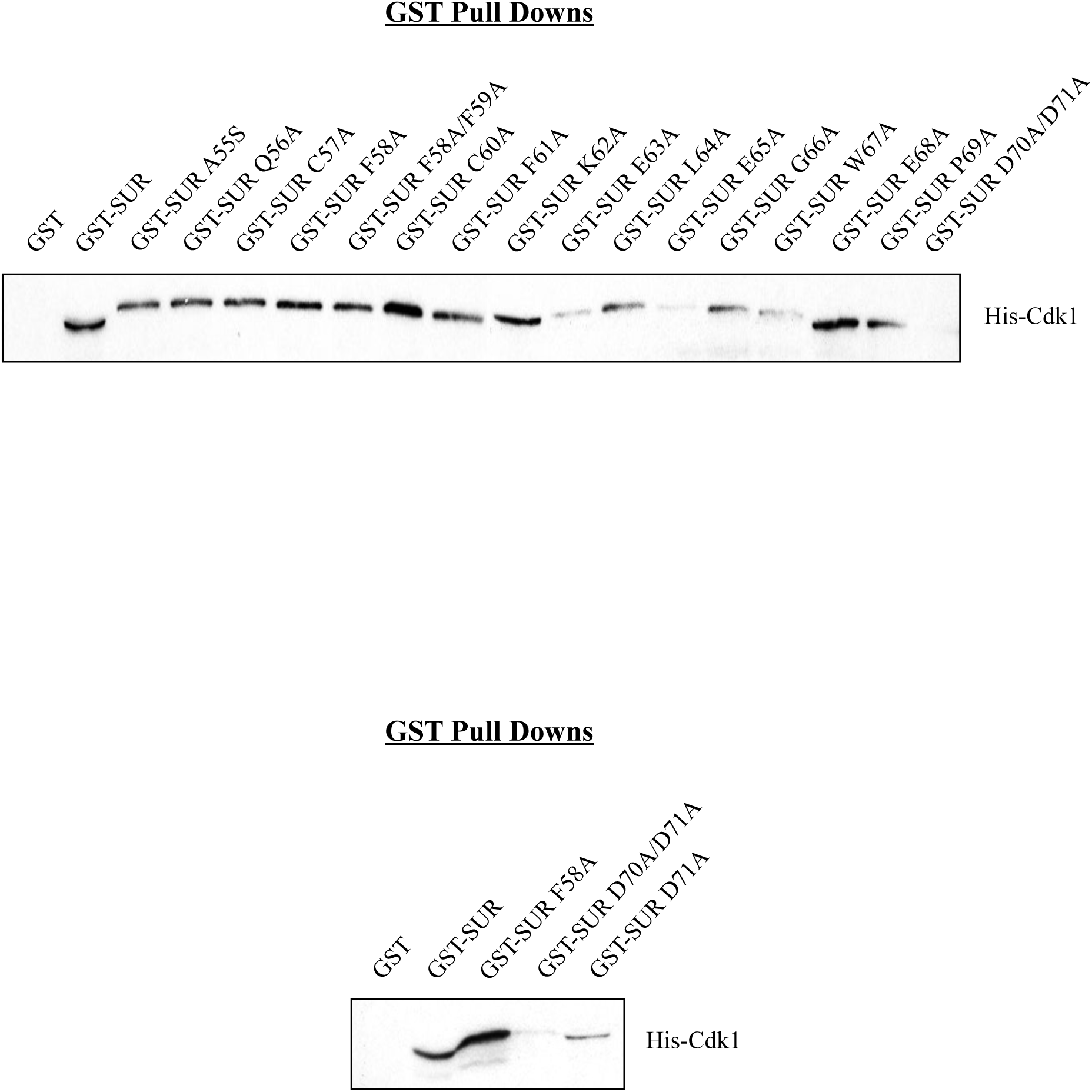
The Survivin Asp70Ala/Asp71Ala (SUR D70A/D71A) double mutant does not bind to Cdk1. Alanine Scanning Mutagenesis of the SUR A55-D70 peptide. The Survivin region spanning the SUR A55-D70 peptide was subjected to alanine mutagenesis, and single alanine GST-Survivin mutants or GST alone were analyzed for their binding to His-Cdk1 by Western blotting (*top*). Binding of SUR D70A/D71A to His-Cdk1 was compared to mutant SUR D71A (*bottom*). An unrelated SUR F58A mutant was used as control.

**FIG. S6.**
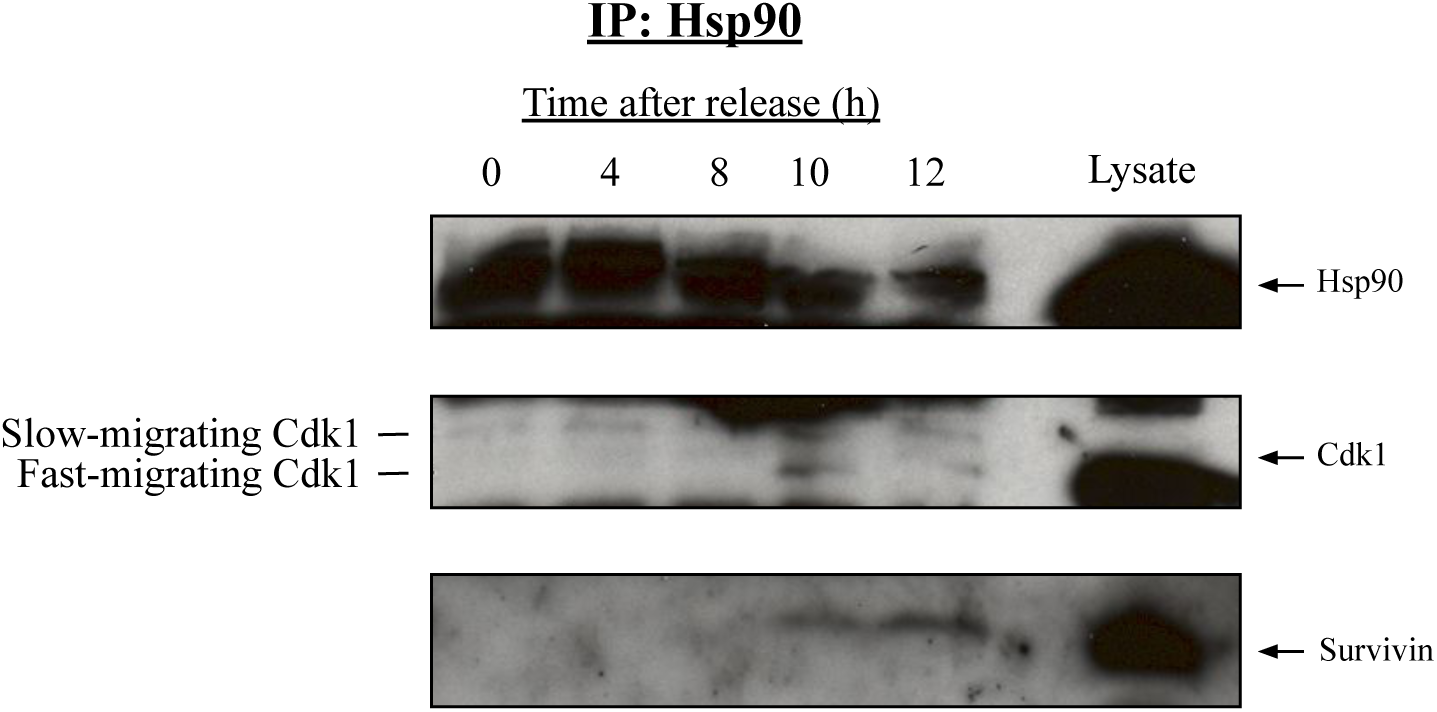
Detection of a Hsp90-Survivin-Cdk1 complex at mitosis. Synchronized HeLa cells were lysed, immunoprecipitated (IP) with an antibody to Hsp90, and analyzed by Western blotting.

## REFERENCES

1. Van der Horst, A. and Lens, S. M. A. (2014). Cell division: control of the chromosomal passenger complex in time and space. Chromosoma 123, 25–42.

2. Lens, S. M. A., Wolthuis, R. M. F., Klompmaker, R., Kauw, J., Agami, R., Brummelkamp, T., Kops, G. and Medema, R. H. (2003). Survivin is required for a sustained spindle checkpoint arrest in response to lack of tension. EMBO J. 22, 2934–2947.

3. Carvalho, A., Carmena, M., Sambade, C., Earnshaw, W. C. and Wheatley, S. P. (2003). Survivin is required for stable checkpoint activation in taxol-treated HeLa cells. J. Cell Sci. 116, 2987–2998.

4. Peter, M., Nakagawa, J., Dorée, M., Labbé, J. C., Nigg, E. A. (1990). In vitro disassembly of the nuclear lamina and M phase-specific phosphorylation of lamins by cdc2 kinase. Cell 61, 591–602.

5. Crncec, A. and Hochegger, H. (2019). Triggering mitosis. FEBS Lett. 593, 2868–2888.

6. Krämer, A., Mailand, N., Lukas, C., Syljuåsen, R. G., Wilkinson, C. J., Nigg, E. A., Bartek, J. and Lukas, J. (2004). Centrosome-associated Chk1 prevents premature activation of cyclin-B-Cdk1 kinase. Nat. Cell Biol. 6, 884–891.

7. Lindqvist, A., Källström, H., Lundgren, A., Barsoum, E. and Rosenthal, C. K. (2005). Cdc25B cooperates with Cdc25A to induce mitosis but has a unique role in activating cyclin B1-Cdk1 at the centrosome. J. Cell Biol. 171, 35–45.

8. Potapova, T. A., Sivakumar, S., Flynn, J. N., Li, R. and Gorbsky, G. J. (2011). Mitotic progression becomes irreversible in prometaphase and collapses when Wee1 and Cdc25 are inhibited. Mol. Biol. Cell 22, 1191–1206.

9. Miake-Lye, R. and Kirschner, M. W. (1985). Induction of early mitotic events in a cell-free system. Cell 41, 165–75.

10. Dessev, G., Iovcheva-Dessev, C., Bischoff, J. R., Beach, D. and Goldman, R. (1991). A complex containing p34cdc2 and cyclin B phosphorylates the nuclear lamin and disassembles nuclei of clam oocytes in vitro. J. Cell Biol. 112, 523–533.

11. Georgatos, S. D., Pyrpasopoulou, A. and Theodoropoulos, P. A. (1997). Nuclear envelope breakdown in mammalian cells involves stepwise lamina disassembly and microtubule-drive deformation of the nuclear membrane. J. Cell Sci. 110, 2129–2140.

12. Beaudouin, J., Gerlich, D., Daigle, N., Eils, R. and Ellenberg. J. (2002). Nuclear envelope breakdown proceeds by microtubule-induced tearing of the lamina. Cell 108, 83–96.

13. Gavet, O. and Pines, J. (2010). Activation of cyclin B1-Cdk1 synchronizes events in the nucleus and the cytoplasm at mitosis. J. Cell Biol. 189, 247–259.

14. Gavet, O. and Pines, J. (2010). Progressive activation of CyclinB1-Cdk1 coordinates entry to mitosis. Dev. Cell 18, 533–543.

15. McCloy, R.A., Rogers, S., Caldon, C. E., Lorca, T., Castro, A. and Burgess, A. (2014). Partial inhibition of Cdk1 in G2 phase overrides the SAC and decouples mitotic events. Cell Cycle 13, 1400–1412.

16. Beltrami, E., Plescia, J., Wilkinson, J. C., Duckett, C. S. and Altieri, D. C. (2004). Acute ablation of survivin uncovers p53-dependent mitotic checkpoint functions and control of mitochondrial apoptosis. J. Biol. Chem. 279, 2077–2084.

17. Cánovas, P. M. and Guadagno, T. M. (2007). Functional analysis of Survivin in spindle assembly in Xenopus egg extracts. J. Cell. Biochem. 100, 217–229.

18. Li, F., Ackermann, E. J., Bennett, C. F., Rothermel, A. L., Plescia, J., Tognin, S., Villa, A., Marchisio, P. C. and Altieri, D. C. (1999). Pleiotropic cell-division defects and apoptosis induced by interference with survivin function. Nat. Cell Biol. 1, 461–466.

19. Giodini, A., Kallio, M. J., Wall, N. R., Gorbsky, G. J., Tognin, S., Marchisio, P. C., Symons, M. and Altieri, D. C. (2002). Regulation of microtubule stability and mitotic progression by survivin. Cancer Res. 62, 2462–2467.

20. Cahu, J., Olichon, A., Hentrich, C., Schek, H., Drinjakovic, J., Zhang, C., Doherty-Kirby, A., Lajoie, G. and Surrey, T. (2008). Phosphorylation by Cdk1 increases the binding of Eg5 to microtubules in vitro and in Xenopus egg extract spindles. PLoS One 3, e3936.

21. Smith, E., Hégarat, N., Vesely, C., Roseboom, I., Larch, C., Streicher, H., Straatman, K., Flynn, H., Skehel, M., Hirota, T., Kuriyama, R. and Hochegger, H. (2011). Differential control of Eg5-dependent centrosome separation by Plk1 and Cdk1. EMBO J. 30, 2233–45.

22. O’Connor, D. S., Grossman, D., Plescia, J., Li, F., Zhang, H., Villa, A., Tognin, S., Marchisio, P. C. and Altieri, D. C. (2000). Regulation of apoptosis at cell division by p34cdc2 phosphorylation of survivin. Proc. Natl. Acad. Sci. U. S. A. 97, 13103–13107.

23. Li, F., Ambrosini, G., Chu, E. Y., Plescia, J., Tognin, S., Marchisio, P. C. and Altieri, D. C. (1998). Control of apoptosis and mitotic spindle checkpoint by survivin. Nature 396, 580–584.

24. Santos, S. D. M., Wollman, R., Meyer, T. and Ferrell Jr., J. E. (2012). Spatial positive feedback at the onset of mitosis. Cell 149, 1500–1513.

25. Pines, J. and Hunter, J. (1991). Human cyclins A and B1 are differentially located in the cell and undergo cell cycle-dependent nuclear transport. J. Cell Biol. 115, 1–17.

26. Tsai, M.-Y., Wang, S., Heidinger, J. M., Shumaker, D. K., Adam, S. A., Goldman, R. D. and Zheng, Y. (2006). A mitotic lamin B matrix induced by RanGTP required for spindle assembly. Science 31, 1887–1893.

27. Borgne, A. and Meijer, L. (1996). Sequential dephosphorylation of p34(cdc2) on Thr-14 and Tyr-15 at the prophase/metaphase transition. J. Biol. Chem. 271, 27847–27854.

28. Brizuela, L, Draetta, G. and Beach, D. (1989). Activation of human CDC2 protein as a histone H1 kinase is associated with complex formation with the p62 subunit. Proc. Natl. Acad. Sci. U. S. A. 86, 4362–4366.

29. Muchmore, S. W., Chen, J., Jakob, C., Zakula, D., Matayoshi, E. D., Wu, W., Zhang, H., Li, F., Ng, S. C. and Altieri, D. C. (2000). Crystal structure and mutagenic analysis of the inhibitor-of-apoptosis protein survivin. Mol. Cell 6, 173–182.

30. Grabarek, J., Amstad, P. and Darzynkiewicz, Z. (2002). Use of fluorescently labeled caspase inhibitors as affinity labels to detect activated caspases. Hum. Cell 15, 1–12.

31. Mazumder, S., Plesca, D. and Almasan, A. (2008). Caspase-3 activation is a critical determinant of genotoxic stress-induced apoptosis. Methods Mol. Biol. 414, 13–21.

32. Chantalat, L., Skoufias, D. A., Kleman, J. P., Jung, B., Dideberg, O. and Margolis, R. L. (2000). Crystal structure of human survivin reveals a bow tie-shaped dimer with two unusual alpha-helical extensions. Mol. Cell 6, 183–189.

33. Keifenheim, D., Sun, X.-M., D’Souza, E., Ohira, M. J., Magner, M., Mayhew, M. B., Marguerat, S., and Rhind, N. (2017). Size-Dependent Expression of the Mitotic Activator Cdc25 Suggests a Mechanism of Size Control in Fission Yeast. Curr. Biol. 27, 1491–1497.

34. Fu, J., Hagan, I. M. and Glover, D. M. (2015). The centrosome and its duplication cycle. Cold Spring Harb. Perspect. Biol. 7, a015800.

35. Norbury, C., Blow, J. and Nurse, P. (1991). Regulatory phosphorylation of the p34cdc2 protein kinase in vertebrates. EMBO J. 10, 3321–3329.

36. Heald, R., McLoughlin, M. and McKeon, F. (1993). Human wee1 maintains mitotic timing by protecting the nucleus from cytoplasmically activated Cdc2 kinase. Cell 74, 463–474.

37. Liu, F., Stanton, J. J., Wu, Z., Piwnica-Worms, H. (1997). The human Myt1 kinase preferentially phosphorylates Cdc2 on threonine 14 and localizes to the endoplasmic reticulum and Golgi complex. Mol. Cell Biol. 17, 571–583.

38. Gottlin, E. B., Xu, X., Epstein, D. M., Burke, S. P., Eckstein, J. W., Ballou, D. P. and Dixon, J. E. (1996). Kinetic analysis of the catalytic domain of human cdc25B. J. Biol. Chem. 271, 27445–27449.

39. Karlsson, C., Katich, S., Hagting, A., Hoffmann, I. and Pines, J. (1999). Cdc25B and Cdc25C differ markedly in their properties as initiators of mitosis. J. Cell. Biol. 146, 573–584.

40. Varmeh-Ziaie, S. and Manfredi, J. J. (2007). The dual specificity phosphatase Cdc25B, but not the closely related Cdc25C, is capable of inhibiting cellular proliferation in a manner dependent upon its catalytic activity. J. Biol. Chem. 282, 24633–24641.

41. Tommasino, M., Accardi, R., Caldeira, S., Dong, W., Malanchi, I., Smet, A. and Zehbe, I. (2003). The role of TP53 in Cervical carcinogenesis. Hum. Mutat. 21, 307–312.

42. Hyun, S.-Y., Rosen, E. M. and Jang, Y.-J. (2012). Novel DNA damage checkpoint in mitosis: Mitotic DNA damage induces re-replication without cell division in various cancer cells. Biochem. Biophys. Res. Commun. 423, 593–5999.

43. Dutertre, S., Cazales, M., Quaranta, M., Froment, C., Trabut, V., Dozier, C., Mirey, G., Bouché, J. P., Theis-Febvre, N., Schmitt, E., Monsarrat, B., Prigent, C. and Ducommun, B. (2004). Phosphorylation of CDC25B by Aurora-A at the centrosome contributes to the G2-M transition. J. Cell Sci. 117, 2523–2531.

44. Nishijima, H., Nishitani, H., Seki, T. and Nishimoto, T. (1997). A dual-specificity phosphatase Cdc25B is an unstable protein and triggers p34(cdc2)/cyclin B activation in hamster BHK21 cells arrested with hydroxyurea. J. Cell Biol. 138, 1105–1116.

45. Gabrielli, B. G., De Souza, C. P., Tonks, I. D., Clark, J. M., Hayward, N. K. and Ellem, K. A. (1996). Cytoplasmic accumulation of cdc25B phosphatase in mitosis triggers centrosomal microtubule nucleation in HeLa cells. J. Cell Sci. 109, 1081–1093.

46. Lammer, C., Wagerer, S., Saffrich, R., Mertens, D., Ansorge, W. and Hoffmann, I. (1998). The cdc25B phosphatase is essential for the G2/M phase transition in human cells. J. Cell Sci. 111, 2445–2453.

47. Baldin, V., Cans, C., Knibiehler, M. and Ducommun, B. (1997). Phosphorylation of human CDC25B phosphatase by CDK1-cyclin A triggers its proteasome-dependent degradation. J. Biol. Chem. 272, 32731–32734.

48. Russell, P. and Nurse, P. (1986). cdc25+ functions as an inducer in the mitotic control of fission yeast. Cell 45, 145–153.

49. Matsusaka, T. and Pines, J. (2004). Chfr acts with the p38 stress kinases to block entry to mitosis in mammalian cells. J. Cell Biol. 166, 507–516.

50. Rieder, C. L. and Cole, R. W. (2000). Microtubule disassembly delays the G2-M transition in vertebrates. Curr. Biol. 10, 1067–1070.

51. Goto, H., Kiyono, T., Tomono, Y., Kawajiri, A., Urano, T., Furukawa, K., Nigg, E. A. and Inagaki, M. (2006). Complex formation of Plk1 and INCENP required for metaphase-anaphase transition. Nat. Cell Biol. 8, 180–187.

52. Zhou, L., Tian, X., Zhu, C., Wang, F. and Higgins, J. M. G. (2014). Polo-like kinase-1 triggers histone phosphorylation by Haspin in mitosis. EMBO Rep. 15, 273–281.

53. Wang, F., Dai, J., Daum, J. R., Niedzialkowska, E., Banerjee, B., Stukenberg P. T., Gorbsky, G. J. and Higgins, J. M. (2010). Histone H3 Thr-3 phosphorylation by Haspin positions Aurora B at centromeres in mitosis. Science 303, 231–235.

54. Wu, Z., Jiang, Q., Clarke, P. R. and Zhang, C. (2013). Phosphorylation of Crm1 by CDK1-cyclin-B promotes Ran-dependent mitotic spindle assembly. J. Cell Sci. 126, 3417–3428.

55. Guo, Li, Mohd, K. S., Ren, H., Xin, G., Jiang, Q., Clarke, P. R. and Zhang, C. (2019). Phosphorylation of importin-α1 by CDK1-cyclin B1 controls mitotic spindle assembly. J. Cell Sci. 132, 232314.

56. O’Connor, D. S., Wall, N. R., Porter, A. C. G. and Altieri, D. C. A p34(cdc2) survival checkpoint in cancer. Cancer Cell 2, 43–54.

57. Matthess, Y., Raab, M., Sanhaji, M., Lavrik, I. N. and Strebhardt, K. (2010). Cdk1/cyclin B1 controls Fas-mediated apoptosis by regulating caspase-8 activity. Mol. Cell Biol. 30, 5726–5740.

58. Allan, L. A. and Clarke, P. R. (2007). Phosphorylation of caspase-9 by CDK1/cyclin B1 protects mitotic cells against apoptosis. Mol. Cell 26, 301–310.

59. Lens, S. M. A., Vader, G. and Medema, R. H. (2006). The case for Survivin as mitotic regulator. Curr. Opin. Cell Biol. 18, 616–622.

60. Altieri, D. C. (2006). The case for survivin as a regulator of microtubule dynamics and cell-death decisions. Curr. Opin. Cell Biol. 18, 609–615.

61. Kannan, N. and Neuwald, A. F. (2005). Did protein kinase regulatory mechanisms evolve through elaboration of a simple structural component? J. Mol. Biol. 351, 956–972.

62. Yeung, W., Ruan, Z. and Kannan, N. (2020). Emerging roles of the αC-β4 loop in protein kinase structure, function, evolution, and disease. IUBMB Life 72, 1189–1202.

63. Chen, H., Ma, J., Li, W., Eliseenkova, A. V., Xu, C., Neubert, T. A., Miller, W. T. and Mohammadi, M. (2007). A molecular brake in the kinase hinge region regulates the activity of receptor tyrosine kinases. Mol. Cell. 27, 717–730.

64. Ruan, Z. and Kannan, N. (2015). Mechanistic Insights into R776H Mediated Activation of Epidermal Growth Factor Receptor Kinase. Biochemistry 54, 4216–4225.

65. Ruan, Z. and Kannan, N. (2018). Altered conformational landscape and dimerization dependency underpins the activation of EGFR by *α* C-*β* 4 loop insertion mutations. Proc. Natl. Acad. Sci. U. S. A. 115, E8162–E8171.

66. Xu, W., Yuan, X., Xiang, Z., Mimnaugh, E., Marcu, M. and Neckers, L. (2005). Surface charge and hydrophobicity determine ErbB2 binding to the Hsp90 chaperone complex. Nat. Struct. Mol. Biol. 12, 120–126.

67. Citri, A., Harari, D., Shohat, G., Ramakrishnan, P., Gan, J., Lavi, S., Eisenstein, M., Kimchi, A., Wallach, D., Pietrokovski, S. and Yarden, Y. (2006). Hsp90 recognizes a common surface on client kinases. J. Biol. Chem. 281, 14361–14369.36.

68. Taipale, M., Krykbaeva, I., Koeva, M., Kayatekin, C., Westover, K. D., Karras, G. I. and Lindquist, S. (2012). Quantitative analysis of HSP90-client interactions reveals principles of substrate recognition. Cell 150, 987–1001.

69. Verba, K. A., Wang, R. Y.-R., Arakawa, A., Liu, Y., Shirouzu, M., Yokoyama, S. and Agard, D. A. (2016). Atomic structure of Hsp90-Cdc37-Cdk4 reveals that Hsp90 traps and stabilizes an unfolded kinase. Science 352, 1542–1547.

70. Plescia, J., Whitney, S., Xia, F., Pennati, M., Zaffaroni, N., Daidone, M. G., Meli, M., Dohi, T., Fortugno, P., Nefedova, Y., Gabrilovich, D. I., Colombo, G. and Altieri, D. C. (2005). Rational design of shepherdin, a novel anticancer agent. Cancer Cell 7, 457–468.

71. Roe, S. M., Ali, M. M. U., Meyer, P., Vaughan, C. K., Panaretou, B., Piper, P. W., Prodromou, C. and Pearl, L. H. (2004). The Mechanism of Hsp90 regulation by the protein kinase-specific cochaperone p50(cdc37). Cell 116, 87–98.

72. Turnbull, E. L., Martin, I. V. and Fantes, P. A. (2006). Activity of Cdc2 and its interaction with the cyclin Cdc13 depend on the molecular chaperone Cdc37 in Schizosaccharomyces pombe. J. Cell Sci. 119, 292–302.

73. García-Morales, P., Carrasco-García, E., Ruiz-Rico, P., Martínez-Mira, R., Menéndez-Gutiérrez, M. P., Ferragut, J. A., Saceda, M. and Martínez-Lacaci, I. (2007). Inhibition of Hsp90 function by ansamycins causes downregulation of cdc2 and cdc25c and G(2)/M arrest in glioblastoma cell lines. Oncogene 26, 7185–7193.

74. Altieri, D. C. (2003). Validating survivin as a cancer therapeutic target. Nat. Rev. Cancer 3, 46–54.

75. Abdelwahab, A. B., El-Sawy, E. R., Hanna, A. G., Bagrel, D. and Kirsch, G. (2022). A Comprehensive Overview of the Developments of Cdc25 Phosphatase Inhibitors. Molecules 27, 2389.

76. Faber, E. B., Sun, L., Tang, J., Roberts, E., Ganeshkumar, S., Wang, N., Rasmussen, D., Majumdar A., Hirsch, L. E., John, K., Yang, A., Khalid, H., Hawkinson, J. E., Levinson, N. M., Chennathukuzhi, V., Harki, D. A., Schönbrunn, E. and Georg, G. I. (2023). Development of allosteric and selective CDK2 inhibitors for contraception with negative cooperativity to cyclin binding. Nat. Commun. 14, 3213.

77. Mesri, M., Wall, N. R., Li, J., Kim, R. W. and Altieri, D. C. (2001). Cancer gene therapy using a survivin mutant adenovirus. J. Clin. Invest. 108, 981–990.

78. Fortugno, P., Beltrami, E., Plescia, J., Fontana, J., Pradhan, D., Marchisio, P. C., Sessa, W. C. and Altieri, D. C. (2003). Regulation of survivin function by Hsp90. Proc. Natl. Acad. Sci. U. S. A. 100, 13791–13796.

